# Twelve Platinum-Standard Reference Genomes Sequences (PSRefSeq) that complete the full range of genetic diversity of Asian rice

**DOI:** 10.1101/2019.12.29.888347

**Authors:** Yong Zhou, Dmytro Chebotarov, Dave Kudrna, Victor Llaca, Seunghee Lee, Shanmugam Rajasekar, Nahed Mohammed, Noor Al-Bader, Chandler Sobel-Sorenson, Praveena Parakkal, Lady Johanna Arbelaez, Natalia Franco, Nickolai Alexandrov, N. Ruaraidh Sackville Hamilton, Hei Leung, Ramil Mauleon, Mathias Lorieux, Andrea Zuccolo, Kenneth McNally, Jianwei Zhang, Rod A. Wing

## Abstract

As the human population grows from 7.8 billion to 10 billion over the next 30 years, breeders must do everything possible to create crops that are highly productive and nutritious, while simultaneously having less of an environmental footprint. Rice will play a critical role in meeting this demand and thus, knowledge of the full repertoire of genetic diversity that exists in germplasm banks across the globe is required. To meet this demand, we describe the generation, validation and preliminary analyses of transposable element and long-range structural variation content of 12 near-gap-free reference genome sequences (RefSeqs) from representatives of 12 of 15 subpopulations of cultivated rice. When combined with 4 existing RefSeqs, that represent the 3 remaining rice subpopulations and the largest admixed population, this collection of 16 Platinum Standard RefSeqs (PSRefSeq) can be used as a pan-genome template to map resequencing data to detect virtually all standing natural variation that exists in the pan-cultivated rice genome.

## Background & Summary

Asian cultivated rice is a staple food for half of the world population. With the planet’s population expected to reach 10 billion by 2050, farmers must increase production by at least 100 million metric tons per year (Seck et al 2012; Merrey et al. 2018). To address this need, future rice cultivars should provide higher yields, be more nutritious, be resilient to multiple abiotic and biotic stresses, and have less of an environmental footprint (Wing et al. 2018; 3K RGP 2014). To achieve this goal, a comprehensive and more in-depth understanding of the full range of genetic diversity of the pan-cultivated rice genome and its wild relatives will be needed (Stein et al. 2018).

With a genome size of ~390 Mb, rice has the smallest genome among the domesticated cereals, making it particularly amenable to genomic studies (Kawahara et al. 2013) and the primary reason why it was the first crop genome to be sequenced 15 years ago (International Rice Genome Sequencing 2005). To better understand the full-range of genetic diversity that is stored in rice germplasm banks around the world, several studies have been conducted using microarrays (Thomson et al. 2017; McNally et al. 2009) and low coverage skim sequencing (Huang et al. 2012; Zhao et al. 2018). In 2018, a detailed analysis of the Illumina resequencing of more than 3,000 diverse rice accessions (a.k.a. 3K-RG), aligned to the *O. sativa* v.g. japonica cv. Nipponbare reference genome sequence (a.k.a. IRGSP RefSeq), showed how the high genetic diversity present in domesticated rice populations provides a solid base for the improvement of rice cultivars (Wang et al. 2018). One key finding from a population structure analysis of this dataset showed that the 3,000 accessions can be subdivided into nine subpopulations, where most accessions from close sub-groups could be associated to geographic origin (Wang et al. 2018).

One critical piece of information missing from these analyses is the fact that single nucleotide polymorphisms (SNPs) and structural variations (SVs) present in subpopulation specific genomic regions have yet to be detected because the 3K-RG data set was only aligned to a single reference genome. Therefore, the next logical step, to capture and understand genetic variation pan-subpopulation-wide is to map the 3K-RG dataset to high-quality reference genomes that represent each of the subpopulations of cultivated Asian rice. At present, only a handful high-quality rice genomes for cultivated rice are publicly available (Kawahara et al. 2013, Zhang et al. 2016a, Zhang et al. 2016b and Stein et al. 2018), thus, there is an immediate need for such a comprehensive resource to be created, which is the subject of this Data Descriptor.

Here we present a reanalysis of the population structure analysis discussed above (Wang et al. 2018) and show that the 3K-RG dataset can be further subdivided into a total of 15 subpopulations. We then present the generation of 12 new and near-gap-free high-quality PacBio long-read reference genomes from representative accessions of the 12 subpopulations of cultivated rice for which no high-quality reference genomes exist. All 12 genomes were assembled with more than 100x genome coverage PacBio long-read sequence data and then validated with Bionano optical maps (Udall and Dawe 2018). The number of contigs covering each of the twelve 12 assemblies, excluding unplaced contigs, ranged from 15 (GOBOL SAIL (BALAM)::IRGC 26624-2) to 104 (IR 64). The contig N50 value for the 12 genome data set ranged from 7.35 Mb to 31.91 Mb. When combined with 4 previously published genomes (i.e. Minghui 63 (MH 63), Zhenshan 97 (ZS 97) (Zhang et al. 2016a, b), N 22 (Stein et al. 2018; updated in 2019) and the IRGSP RefSeq (Kawahara et al. 2013)), this 16 genome dataset can be used to represent the K=15 population/admixture structure of cultivated Asian rice.

## Methods

### Ethics statement

This work was approved by the University of Arizona (UA), the King Abdullah University of Science and Technology (KAUST), Huazhong Agricultural University (HZAU), the International Rice Research Institute (IRRI) and the International Center for Tropical Agriculture (CIAT). All methods used in this study were carried out following approved guidelines.

### Population structure

We extracted 30 subsets of 100,000 randomly chosen SNPs out of the 3K-RG Core SNP set v4 (996,009 SNPs, available at https://snp-seek.irri.org/_download.zul). For each subset, we ran ADMIXTURE (Alexander et al. 2009) with the number of ancestral groups K ranging from 5 to 15. We then aligned the resulting Q matrices using CLUMPP software (Jakobsson and Rosenberg 2007). Since different runs at a given value of K often give rise to different refinements (splits) of the lower level grouping, we first clustered the runs for each K according to similarity of Q matrices using hierarchical clustering, thus obtaining several clusters of runs for each K. We discarded one-element clusters (outlier runs), and averaged the Q matrices within each remaining cluster. Figure S1 shows the admixture proportions taken from the averaged Q matrices of the final clusters for K=9 to 15. The columns of these averaged Q matrices, representing admixture proportions for groups discovered in different runs, were then used to define the “K15” grouping. At K=9, 12, and 13, the Q matrices converged to two different modes according to whether XI-1A or GJ-trop is split (these are labeled as K=9.1, 12.1 and 13.1).

The group membership for each sample was defined by applying the threshold of 0.65 to admixture components. Samples with no admixture components exceeding 0.65 were classified as follows. If the sum of components for subpopulations within the major groups cA (*circum*-Aus), XI (*Xian*-indica), and GJ (*Geng*-japonica) was ≥ 0.65, the samples were classified as cA-adm (admixed within cA), XI-adm (within XI) or GJ-adm (within GJ), respectively, and the remaining samples were deemed ‘fully’ admixed. The newly defined groups were mostly either aligned with the previous K=9 grouping, or refined those groups, and they were named accordingly (e.g. XI-1B1 and XI-1B2 are new subgroups within XI-1B).

The phenogram shown in Figure 1 was constructed with DARwin v6 (http://darwin.cirad.fr/, unweighted Neighbor-joining) using the identity by state (IBS) distance matrix from Plink on the 4.8M Filtered SNP set (available at https://snp-seek.irri.org/_download.zul). Colors were assigned to subpopulations based on K15 Admixture results. One entry, MH 63 (XI-adm) represents the admixed types among the XI group.

**Figure 1.**
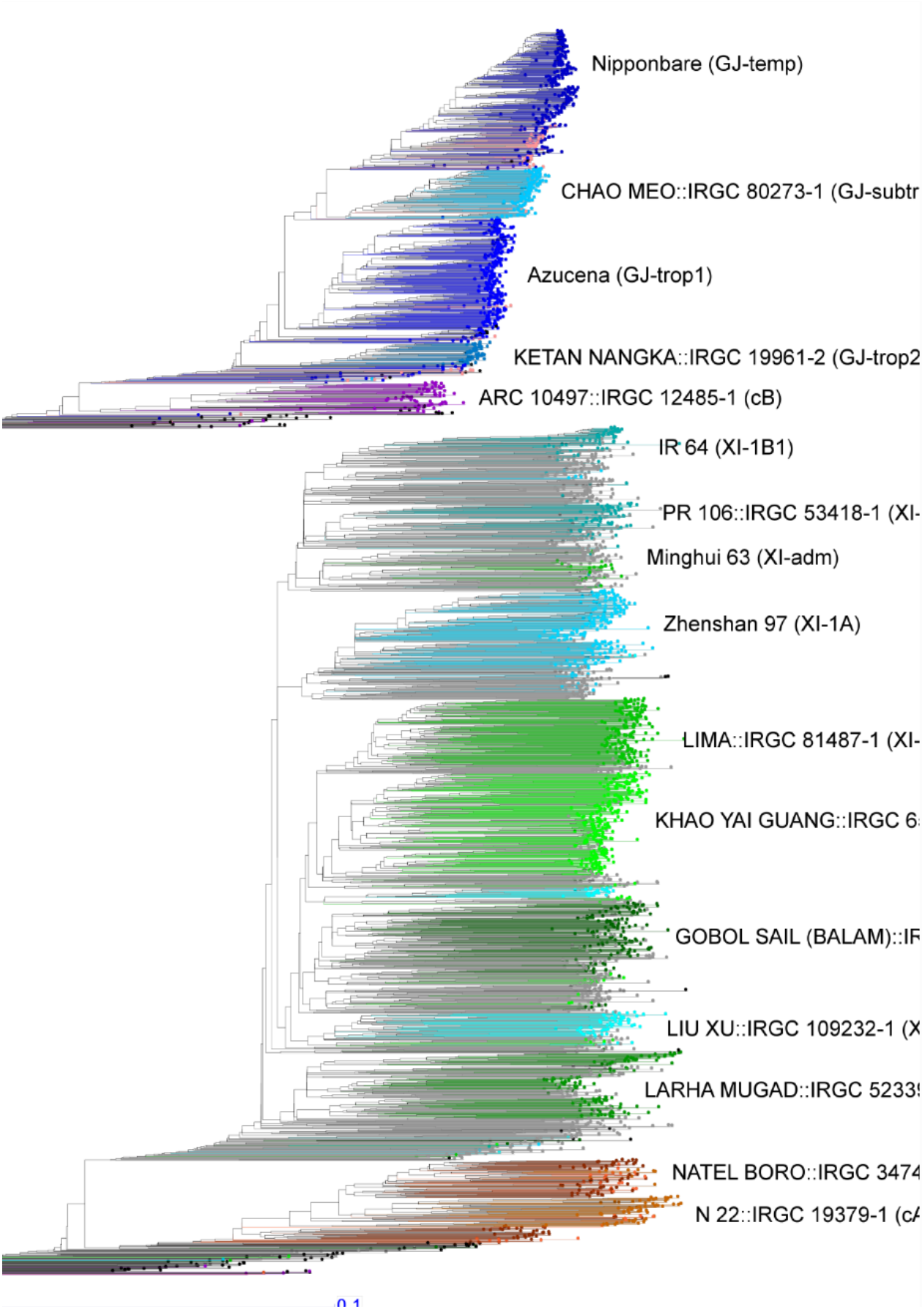
Phylogenetic tree with the accession selected for PSRefSeq sequencing for each of the K=15 subpopulations and a single admixture group. Groups are colored according to the assignment from Admixture analysis. The subpopulation designation is in parentheses following the name.

### Sample selection, collection and nucleic acid preparation

To select accessions to represent the 12 subpopulations of Asian rice that lack high-quality reference genome assemblies, the following strategy was employed. The IBS distance matrix was used for a principlal component analysis (PCA) analysis in R to generate 5 component axes. Then, for each of the 12 subpopulations, i.e. *circum*-Aus2 = cA2, *circum*-Basmati = cB, *Geng*-japonica (GJ) subtropical (GJ-subtrp), tropical1 (GJ-trop1) and tropical2 (GJ-trop2), and *Xian*-indica (XI) subpopulations XI-1B1, XI-1B2, XI-2A, XI-2B, XI-3A, XI-3B1, XI-3B2, the centroid of each group in the space spanned by first 5 principal components was determined from the eigenvectors, and the entry closest to the centroid for which seed was available was chosen as the representative for that subpopulation (Table 1).

**Table 1.** Sample collection information for 12 *Oryza sativa* accessions.

| Variety Name | Genetic Stock ID | Country Origin | 12 subpopulations |
| --- | --- | --- | --- |
| CHAO MEO::IRGC 80273-1 | IRGC 132278 | Lao PDR | GJ-subtrp |
| Azucena | I1A41685 | Philippines | GJ-trop1 |
| KETAN NANGKA::IRGC 19961-2 | IRGC 128077 | Indonesia | GJ-trop2 |
| ARC 10497::IRGC 12485-1 | IRGC 117425 | India | cB |
| IR 64 | I1A42114 | Philippines | XI-1B1 |
| PR 106::IRGC 53418-1 | IRGC 127742 | India | XI-1B2 |
| LIMA::IRGC 81487-1 | IRGC 127564 | Indonesia | XI-3A |
| KHAO YAI GUANG::IRGC 65972-1 | IRGC 127518 | Thailand | XI-3B1 |
| GOBOL SAIL (BALAM)::IRGC 26624-2 | IRGC 132424 | Bangladesh | XI-2A |
| LIU XU::IRGC 109232-1 | IRGC 125827 | China | XI-3B2 |
| LARHA MUGAD::IRGC 52339-1 | IRGC 125619 | India | XI-2B |
| NATEL BORO::IRGC 34749-1 | IRGC 127652 | Bangladesh | cA2 |

Subpopulations: GJ = *Geng-japonica* where trop = tropical, subtrp = subtropical; cB = *circum-Basmati*; XI = *Xian-indica*; cA = *circum-Aus*

Single seed decent (SSD) seed from IR 64 and Azucena were obtained from the Rice Genetics and Genomics Laboratory, CIAT, in Cali, Colombia, and SSD seed from the remaining 10 accessions (Table 1) were obtained from the International Rice Genebank, maintained by IRRI, Los Baños, Philippines. All seed were sown in potting soil and grown under standard greenhouse conditions at UA, Tucson, USA for 6 weeks at which point they were dark treated for 48-hours to reduce starch accumulation. Approximately 20-50 grams of young leaf tissue was then harvested from each accession and immediately flash frozen in liquid nitrogen before being stored at −80°C prior to DNA extraction. High molecular weight genomic DNA was isolated using a modified CTAB procedure as previously described (Porebski et al. 1997). The quality of each extraction was checked by pulsed-field electrophoresis (CHEF) on 1% agarose gels for size and restriction enzyme digestibility, and quantified by Qubit fluorometry (Thermo Fisher Scientific, Waltham, MA).

### Library construction and sequencing

Genomic DNA from all 12 accessions were sequenced using the PacBio single-molecule real-time (SMRT) platform, and the Illumina platform for genome size estimations and sequence polishing. High molecular weight (HMW) DNA from each accession was gently sheared into large fragments (*i.e.* 30Kb - 60Kb) using 26-gauge needles and then end-repaired according to manufacturer’s instructions (Pacific Biosciences). Briefly, using a SMRTbell Express Template Prep Kit, blunt hairpins and sequencing adaptors were ligated to HMW DNA fragments, and DNA sequencing polymerases were bound to the SMRTbell templates. Size selection of large fragments (above 15Kb) was performed using a BluPippin electrophoresis system (Sage Science). The libraries were quantified using a Qubit Fluorometer (Invitrogen, USA) and the insert mode size was determined using an Agilent fragment analyzer system with sizes ranging between 30Kb - 40Kb. The libraries then were sequenced using SMRT Cell 1M chemistry version 3.0 on a PacBio Sequel instrument. The number of long-reads generated per accession ranged from 2.01 million (LIMA::IRGC 81487-1) to 5.40 million (Azucena). The distribution of subreads is shown in Figure S2 and the average lengths ranged from 10.58 Kb (Azucena) to 20.61 Kb (LIMA::IRGC 81487-1) (Table 2). According to the estimated genome size of the IRGSP RefSeq, the average PacBio sequence coverage for each accession varied from 103x (LIMA::IRGC 81487-1) to 149x (IR 64) (Table 2).

**Table 2.** Sequencing platforms used and data statistics for the 12 *Oryza sativa* genomes.

| Variety Name | Sequencing platform | Raw data (Gb) | Depth | Number of subreads (M) | Mean subread length (Kb) |
| --- | --- | --- | --- | --- | --- |
| CHAO MEO::IRGC 80273-1 | PacBio Sequel | 49.1 | 123X | 4.26 | 11.526 |
| Azucena | PacBio Sequel | 57.1 | 143X | 5.40 | 10.581 |
| KETAN NANGKA::IRGC 19961-2 | PacBio Sequel | 49.8 | 125X | 2.78 | 17.876 |
| ARC 10497::IRGC 12485-1 | PacBio Sequel | 44.7 | 112X | 4.06 | 11.026 |
| IR 64 | PacBio Sequel | 59.7 | 149X | 5.24 | 11.393 |
| PR 106::IRGC 53418-1 | PacBio Sequel | 42.2 | 105X | 2.08 | 20.317 |
| LIMA::IRGC 81487-1 | PacBio Sequel | 41.4 | 103X | 2.01 | 20.612 |
| KHAO YAI GUANG::IRGC 65972-1 | PacBio Sequel | 42.5 | 106X | 2.37 | 17.954 |
| GOBOL SAIL (BALAM)::IRGC 26624-2 | PacBio Sequel | 42.2 | 105X | 2.13 | 19.777 |
| LIU XU::IRGC 109232-1 | PacBio Sequel | 55.3 | 138X | 3.66 | 15.109 |
| LARHA MUGAD::IRGC 52339-1 | PacBio Sequel | 45.1 | 113X | 3.22 | 14.011 |
| NATEL BORO::IRGC 34749-1 | PacBio Sequel | 44.4 | 111X | 2.74 | 16.2 |

For Illumina short-read sequencing, HMW DNA from each accession was sheared to between 250-1000bp, followed by library construction targeting 350bp inserts following standard Illumina protocols (San Diego, CA, USA). Each library was 2 × 150bp paired-end sequenced using an Illumina X-ten platform. Low-quality bases and paired reads with Illumina adaptor sequences were removed using *Trimmomatic* (Bolger et al. 2014). Quality control for each library data set was carried out with *FastQC* (Brown et al. 2017). Finally, between 36.52-Gb and 51.05-Gb of clean data from each accession was generated and used for genome size estimation (Table S1) by Kmer analysis (Figure S3) and the Genome Characteristics Estimation (GCE) program (Liu et al. 2013).

### Bionano optical genome maps

Bionano optical maps for each accession were generated as previously described (Ou et al. 2019), except that ultra-HMW DNA isolation, from approximately 4g of flash-frozen dark-treated (48 hour) leaf tissue per accession, was performed according to a modified version of the protocol described by Luo and Wing (Luo and Wing, 2003). Prior to labeling, agarose plugs were digested with agarase and the starch and debris removed by short rounds of centrifugation at 13,000 X g. DNA samples were further purified and concentrated by drop dialysis against TE Buffer. Data processing, optical map assembly, hybrid scaffold construction and visualization were performed using the Bionano Solve (Version 3.4) and Bionano Access (v12.5.0) software packages (https://bionanogenomics.com/).

### *De novo* genome assembly

Genome assembly for each of the 12 genomes followed a five-step procedure as shown in (Figure 2):

Step 1: PacBio subreads were assembled *de novo* into contigs using three genome assembly programs: FALCON (Chin et al. 2016), MECAT2 (Xiao et al. 2017) and Canu1.5 (Koren et al. 2017). The number of *de novo* assembled contigs obtained varied from 51 (e.g. NATEL BORO::IRGC 34749-1 and KETAN NANGKA::IRGC 19961-2) to 1,473 (CHAO MEO::IRGC 80273-1) for the 12 genomes (Table S2).
Step 2: Genome Puzzle Master (GPM) software (Zhang et al. 2016c) was used to merge the *de novo* assembled contigs from the three assemblers, using the high-quality *O. sativa* vg. indica cv. Minghui 63 reference genome MH63RS2 (Zhang et al. 2016a,b) as a guide. GPM is a semi-automated pipeline that is used to integrate logical relationship data (*i.e.* contigs from three assemblers for each accession) based on a reference guide. Contigs were merged in the ‘assemblyRun’ step, with default parameters (minOverlapSeqToSeq was set at 1 Kb and identitySeqToSeq was set at 99%). Redundant overlapping sequences were also removed for each assembled contig. In addition, we gave contiguous contigs a higher priority than ones with gaps to be retained in each assembly. After manual checking, editing, and redundancy removal, the number of contigs in each assembly ranged from 26 (NATEL BORO::IRGC 34749-1) to 588 (LIU XU::IRGC 109232-1) (Table S3).
Step 3: The sequence quality of each contig was then improved by “sequence polishing”: twice with PacBio long reads and once with Illumina short reads. Briefly, PacBio subreads were aligned to GPM edited contigs using the software *blasr* (Chaisson and Tesler 2012). All default parameters were used, except minimum align length, which was set to 500-bp. Secondly, the tool *arrow* as implemented in SMRTlink6.0 (Pacific Biosciences of California, Inc) was used for polishing the GPM edited contigs. The *bwa-mem* program (Li 2013) was then used for mapping short Illumina reads onto assembled contigs, and the tool *pilon* (Walker et al. 2014) was used for a final polishing step with default settings.
Step 4: The polished contigs for each accession were arranged into pseudomolecules using *GPM*, using MH63RS2 (Zhang et al. 2016a,b) as the reference guide. The program *blastn* (Altschul et al. 1997) with a minimum alignment length of 1 Kb and an e-value < 1e^−5^ as the threshold was used to align the corrected contigs to the reference guide. In doing so, the corrected contigs were assigned to chromosomes, as well as ordered and orientated in the GPM assembly viewer function. The number of contigs after step 4 ranged from a minimum of 15 contigs (GOBOL SAIL (BALAM)::IRGC 26624-2) to a maximum of 104 contigs (IR64) (Table 3). The assembly size for the 12 accessions ranged from 376.86 Mb (CHAO MEO::IRGC 80273-1) to 393.74 Mb (KHAO YAI GUANG::IRGC 65972-1) (Table 3) and the length of individual chromosome varied from 23.06 Mb (chromosome 9 of CHAO MEO::IRGC 80273-1) to 44.96 Mb (chromosome 1 of LIMA::IRGC 81487-1) (Table S4). The average N50 value was 23.10 Mb, with the highest and the lowest values being 30.91 Mb (LIU XU::IRGC 109232-1) and 7.35 Mb (IR 64), respectively. The average number of gaps among the 12 new genome assemblies was 18, with 8 assemblies containing less than 10 gaps (Table 3).
Step 5: To independently validate our assemblies, we generated and compared Bionano optical maps to each assembly. In total, 17 (Azucena) to 56 (LIU XU::IRGC 109232-1) Bionano optical maps were constructed for all 12 rice accessions, which yielded contig N50 values of between 22.75 Mb (CHAO MEO::IRGC 80273-1) to 31.45 Mb (KHAO YAI GUANG::IRGC 65972-1) (Table S5). As shown in Figure 3 and Figure S4, the chromosomes and/or chromosome arms of all 12 *de novo* assemblies were highly supported by these ultra-long optical maps. Although rare, a few discrepancies between the optical maps and genome assemblies can be seen and are likely due to small errors and chimeras that can be produced through both the optical mapping and sequence assembly pipelines (Udall and Dawe 2018).

**Figure 2.**
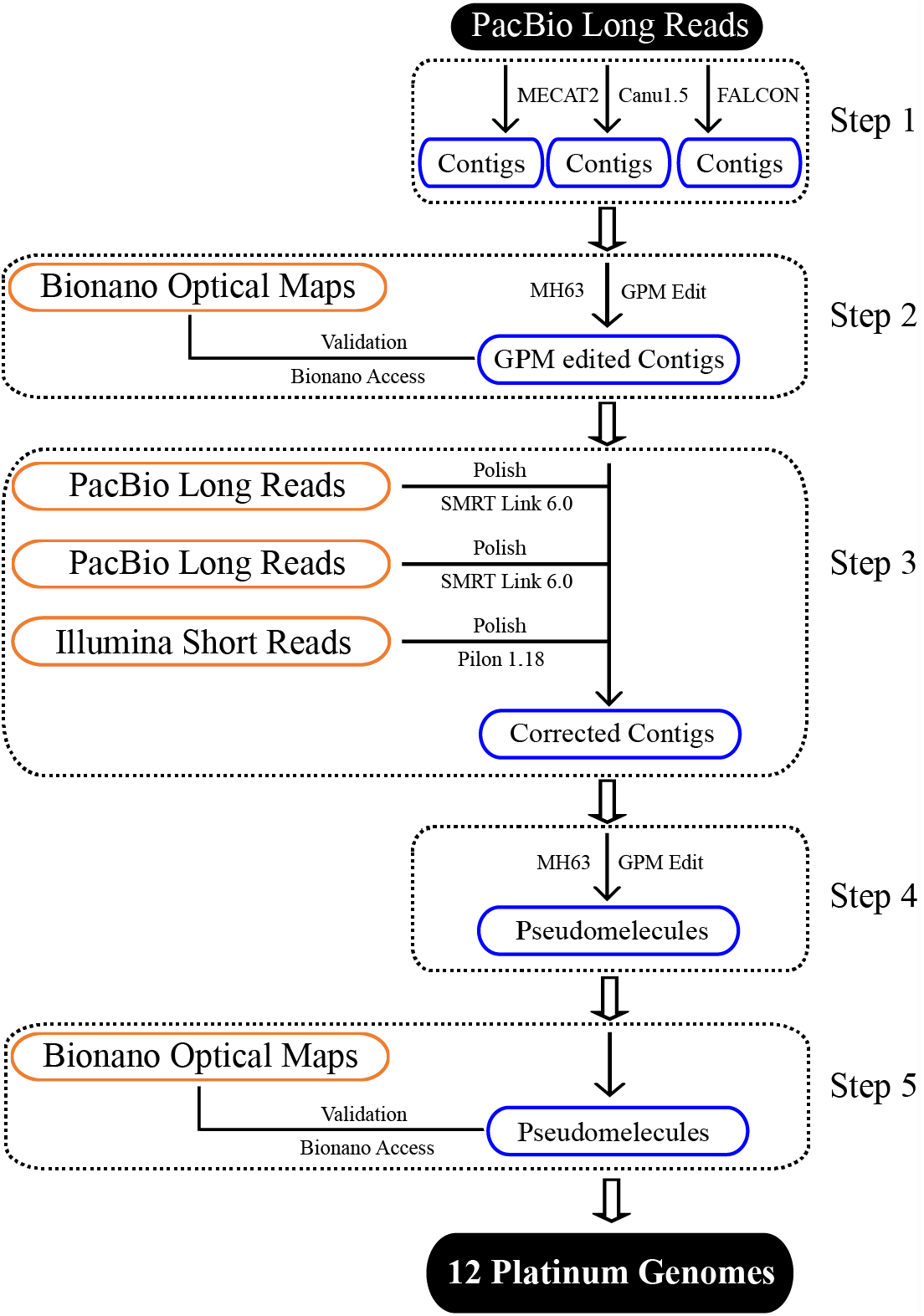
Genome assembly and validation pipeline.

**Table 3.** *de novo* assembly, BUSCO evaluation and accession numbers in GenBank of the 12 *Oryza sativa* genomes.

| Variety Name | BioProject | BioSample | Genome size (bp) | # Contigs | Contig N50 (Mb) | # Gaps | Scaffold N50 (Mb) | BUSCO | Adjust BUSCO | Genome Accession | SRP | Supplementary Files (Bionano optical map) |
| --- | --- | --- | --- | --- | --- | --- | --- | --- | --- | --- | --- | --- |
| CHAO MEO::IRGC 80273-1 | PRJNA565484 | SAMN12748601 | 376,856,903 | 55 | 11.02 | 43 | 30.35 | 97.60% | 98.49% | VYIH000000000 | SRP226088 | SUPPF_0000003210 |
| Azucena | PRJNA424001 | SAMN08217222 | 379,627,553 | 28 | 22.94 | 16 | 30.95 | 97.80% | 98.69% | PKQC000000000 | SRP227255 | SUPPF_0000003212 |
| KETAN NANGKA::IRGC 19961-2 | PRJNA564615 | SAMN12718029 | 380,759,091 | 21 | 22.68 | 9 | 30.70 | 98.00% | 98.89% | VYIC000000000 | SRP226080 | SUPPF_0000003204 |
| ARC 10497::IRGC 12485-1 | PRJNA565479 | SAMN12748569 | 378,463,869 | 40 | 17.92 | 28 | 30.57 | 98.40% | 99.30% | VYID000000000 | SRP226093 | SUPPF_0000003206 |
| IR 64 | PRJNA509165 | SAMN10564385 | 386,698,898 | 104 | 7.35 | 92 | 31.22 | 95.70% | 96.57% | RWKJ000000000 | SRP227298 | SUPPF_0000003213 |
| PR 106::IRGC 53418-1 | PRJNA563359 | SAMN12672924 | 391,176,105 | 16 | 27.05 | 4 | 32.03 | 96.60% | 97.48% | VYIB000000000 | SRP226078 | SUPPF_0000003202 |
| LIMA::IRGC 81487-1 | PRJNA564572 | SAMN12715984 | 392,625,308 | 17 | 27.37 | 5 | 32.42 | 98.50% | 99.40% | VXJH000000000 | SRP226079 | SUPPF_0000003203 |
| KHAO YAI GUANG::IRGC 65972-1 | PRJNA565481 | SAMN12748590 | 393,737,720 | 19 | 21.82 | 7 | 32.08 | 98.60% | 99.50% | VYIF000000000 | SRP226086 | SUPPF_0000003208 |
| GOBOL SAIL (BALAM)::IRGC 26624-2 | PRJNA564763 | SAMN12721963 | 391,772,995 | 15 | 29.60 | 3 | 31.75 | 97.90% | 98.79% | VXJI000000000 | SRP226082 | SUPPF_0000003205 |
| LIU XU::IRGC 109232-1 | PRJNA577228 | SAMN13021815 | 392,033,263 | 17 | 30.91 | 5 | 32.30 | 98.40% | 99.30% | WGGU000000000 | SRP226085 | SUPPF_0000003211 |
| LARHA MUGAD::IRGC 52339-1 | PRJNA565480 | SAMN12748589 | 390,195,943 | 16 | 30.75 | 4 | 32.10 | 98.60% | 99.50% | VYIE000000000 | SRP226084 | SUPPF_0000003207 |
| NATEL BORO::IRGC 34749-1 | PRJNA565483 | SAMN12748600 | 383,720,936 | 16 | 27.83 | 4 | 31.31 | 98.10% | 98.99% | VYIG000000000 | SRP226087 | SUPPF_0000003209 |

**Figure 3.**
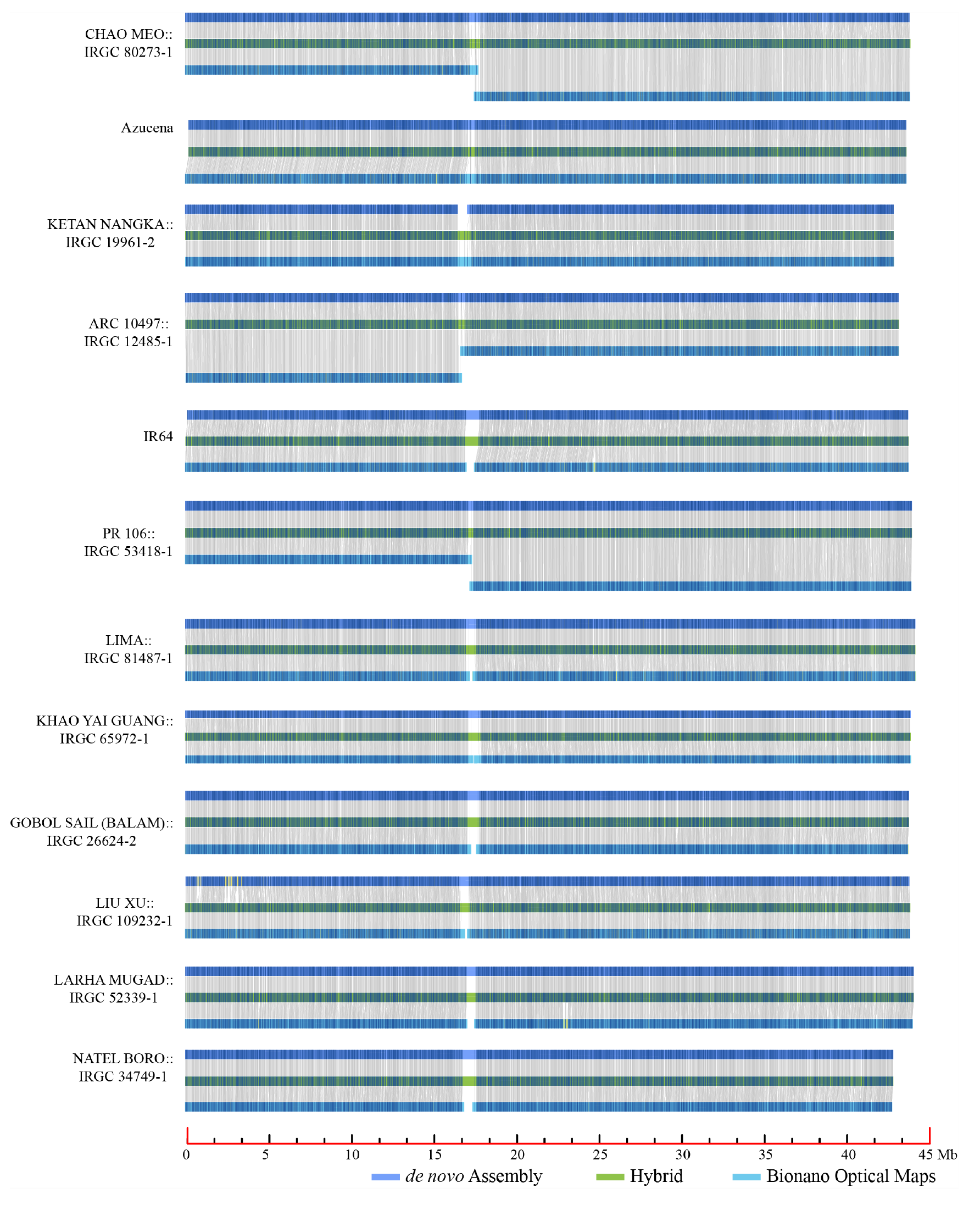
Bionano optical map validation of chromosome 1 for 12 *de novo* assemblies.

Following these five steps, we were able to produce 12 near-gap-free *Oryza sativa* platinum standard reference genome sequences (PSRefSeqs) that represent 12 of 15 subpopulations of cultivated Asian rice.

### BUSCO evaluation

The Benchmarking Universal Single-Copy Orthologs (BUSCO3.0) software package (Simao et al. 2015) was employed to evaluate the gene space completeness of the 12 genome assemblies. These genomes captured, on average, 97.9% of the BUSCO reference gene set, with a minimum of 95.7% (IR64) and a maximum of 98.6% (LARHA MUGAD::IRGC 52339-1 and KHAO YAI GUANG::IRGC 65972-1) (Table 3).

Of note, when performing this analysis, we observed that on average 30 out of the 1,440 conserved BUSCO genes tested (https://www.orthodb.org/v9/index.html) were missing from each new assembly, 16 of which were not present in all 12, plus the IRGSP, ZS 97, MH 63 and N 22 RefSeqs (Figure S5). This result suggested that these 16 “conserved” genes may not exist in rice, or other cereal genomes, thereby artificially reducing the BUSCO gene space scores for our 12 assemblies. To test this hypothesis, we searched for all 16 genes missing in maize, which diverged from rice about 50 million years ago (MYA) (Wolfe et al., 1989, Gale et al., 1998 and Guo et al., 2019). We found that 13 of the 16 genes in question could not be found in 3 high-quality recently published maize genome assemblies (Figure S5) and therefore, concluded that 13 of the 16 “conserved” genes in the BUSCO database are not present in cereals, and should be excluded from our gene space analysis. Taking this into account, we recalculated the BUSCO gene space content for each of 12 assemblies and found that 10 of 12 assemblies captured more than 98% of the BUSCO gene set (Table 3).

### Transposable element (TE) prediction

To determine the pan-transposable element content of cultivated Asian rice we analyzed the 12 new reference genomes, presented here, along with the MH 63, ZS 97, N 22 PacBio reference genomes. In addition, we also included a reanalysis of the IRGSP RefSeq as it is conventionally considered the standard rice genome for which all comparisons are conducted. This 16 genome data set was used to represent the K=15 population structure of cultivated Asian rice.

A search for sequences similar to TEs was carried out using RepeatMasker (Smit AFA et al, 2013) run under default parameters with the exception of the options: `-no_is -nolow`. RepeatMasker was run using the library “rice 7.0.0.liban”, which is an updated in-house version of the publicly available MSU_6.9.5 library (Ou et al. 2019), retrieved from https://github.com/oushujun/EDTA/blob/master/database/Rice_MSU7.fasta.std6.9.5.out. The average TE content of this 16 genome data set was 47.66% with a minimum value of 46.07% in IRGSP RefSeq and a maximum of 48.27% in KHAO YAI GUANG::IRGC 65972-1 (Table 4). The major contribution to this fraction was composed of long terminal repeat retrotransposons (LTR-RTs, min: 23.55%, max: 27.27% and average: 25.96%) followed by DNA-TEs (min:14.87%, max, 16.18% and average: 15.26%). Long interspersed nuclear elements (LINEs) and short interspersed nuclear elements (SINEs) were identified as on average 1.43% and 0.39% of the 16 genomes, respectively.

**Table 4.** Abundance of the major TE classes in the 16 *Oryza sativa* genomes.

| Variety Name | TOTAL | LTR-RT | LINEs | SINEs | DNA_TEs | Unclassified |
| --- | --- | --- | --- | --- | --- | --- |
| NIPPONBARE | 46.07 | 23.55 | 1.52 | 0.41 | 16.18 | 4.41 |
| CHAO MEO::IRGC 80273-1 | 46.25 | 24.00 | 1.46 | 0.40 | 15.59 | 4.80 |
| Azucena | 47.07 | 24.48 | 1.47 | 0.40 | 15.82 | 4.89 |
| KETAN NANGKA::IRGC 19961-2 | 46.99 | 24.87 | 1.47 | 0.40 | 15.72 | 4.53 |
| ARC 10497::IRGC 12485-1 | 46.95 | 24.74 | 1.48 | 0.40 | 15.68 | 4.65 |
| IR 64 | 47.87 | 26.82 | 1.42 | 0.40 | 14.97 | 4.26 |
| PR 106::IRGC 53418-1 | 47.95 | 26.82 | 1.41 | 0.39 | 15.05 | 4.28 |
| Minghui 63 | 47.97 | 26.61 | 1.44 | 0.4 | 15.3 | 4.22 |
| Zhenshan 97 | 47.95 | 26.79 | 1.42 | 0.39 | 15.19 | 4.16 |
| LIMA::IRGC 81487-1 | 48.04 | 26.87 | 1.40 | 0.39 | 15.01 | 4.37 |
| KHAO YAI GUANG::IRGC 65972-1 | 48.27 | 27.27 | 1.40 | 0.39 | 14.87 | 4.34 |
| GOBOL SAIL (BALAM)::IRGC 26624-2 | 48.15 | 26.99 | 1.40 | 0.39 | 14.99 | 4.38 |
| LIU XU::IRGC 109232-1 | 46.92 | 27.06 | 1.26 | 0.32 | 14.31 | 3.97 |
| LARHA MUGAD::IRGC 52339-1 | 48.05 | 26.74 | 1.41 | 0.39 | 15.09 | 4.42 |
| NATEL BORO::IRGC 34749-1 | 47.33 | 25.75 | 1.42 | 0.40 | 15.12 | 4.64 |
| N 22::IRGC 19379-1 | 47.79 | 25.95 | 1.44 | 0.39 | 15.20 | 4.81 |

### Structural Variants

Each genome assembly (n=16), as described above, was fragmented using the EMBOSS tool *splitter* (Rice et al. 2000) to create a 10x genome equivalent redundant set of 50kb reads. These reads were then mapped onto every other genome assembly using the tool *NGMLR* (Sedlazeck et al. 2018). Finally, the software *SVIM* (Heller and Vingron 2019) was run under default parameters to parse the mapping output. Only insertions, deletions and tandem duplications up to a maximum length of 25 Kb were considered in this analysis.

The results of this analysis identified several thousand insertions and deletions whenever an assembly was compared to any other. Greater variability was found between varieties belonging to different major groups (e.g. *Geng*-japonica vs. *Xian*-indica) than occurred between those within these groups. The amount of genome sequences with structural variation between any two varieties ranged from 17.57 Mb to 41.54 Mb for those belonging to the indica (XI, *Xian*-indica) varietal group (avg: 31.75 Mb) and from 18.55 Mb to 23.07 Mb (avg: 21.00 Mb) for those in the japonica (GJ, *Geng*-japonica) varietal group. When all 16 genomes are considered together, the range is between 17.57 Mb and 41.54 Mb, with an average value of 33.70 Mb (Table S6). The total unshared fraction collected out of all pairwise comparisons was composed for 89.89% by TE related sequences.

### Data Records

Data for all 12 genome shotgun sequencing projects have been deposited in Genbank (https://www.ncbi.nlm.nih.gov/), including PacBio raw data, Illumina raw data, Bionano optical maps and the twelve PSRefSeqs. The BioProjects, BioSamples, Genome assemblies, Sequence Read Archives (SRA) accession and supplementary files (Bionano optical maps) of 12 genomes are listed in Table 3.

### Technical Validation

#### DNA sample quality

DNA quality was checked by pulsed-field gel electrophoresis for size and restriction enzyme digestibility. Nucleic acid concentrations were quantified by Qubit fluorometry (Thermo Fisher Scientific, Waltham, MA).

#### Illumina libraries

Illumina libraries were quantified by qPCR using the KAPA Library Quantification Kit for Illumina Libraries (KapaBiosystems, Wilmington, MA, USA), and library profiles were evaluated with an Agilent 2100 Bioanalyzer (Agilent Technologies, Santa Clara, CA, USA).

#### Gene Space Completeness

Benchmarking Universal Single-Copy Orthologs (BUSCO3.0) was executed using the embryophyta_odb9.tar.gz database to assess the gene space of each genome, minus 13 genes that do not appear to exist in the cereal genomes tested (Figure S5).

#### Assembly accuracy

Bionano optical maps were generated and used to validate all 12 genome assemblies.

This paper is the first release of 12 PSRefSeqs, optical maps and all associated raw data for the accessions listed in Table 3.

### Code Availability

The population re-analysis of 3K-RG dataset and 12 genome assemblies were obtained using several publicly available software packages. To allow researchers to precisely repeat any steps, the settings and the parameters used are provided below:

#### Population structure

ADMIXTURE (Alexander et al. 2009) was run with default options. The R scripts for further population structure analysis, including setting up CLUMPP files, can be found in Github repository https://github.com/dchebotarov/Q-aggr.

#### Genome size estimation

The K-mer and GCE program (Liu et al. 2013) were employed for genome size estimation. Command line:

~~~
kmer_freq_hash -k (13-17) -l genome.list -a 10 -d 10 -t 8 -
i 400000000 -o 0 -p genom_kmer(13-17) &> genome_kmer(13-
17)_freq.log, and gce -f genom _kmer(13-17).freq.stat -c
$peak $g #amount -m 1 -D 8 -b 1 -H 1 > genome.table 2>
genom_kmer(13-17).log
~~~

#### Genome assembly

1. *MECAT2*: all parameters were set to the defaults. Command line:

~~~
mecat.pl config_file.txt, mecat.pl correct config_file.txt
and mecat.pl assemble config_file.txt
~~~
2. *Canu1.5*: all parameters were set to the defaults. Command line:

~~~
canu -d canu -p canu genomeSize=400m -pacbio-raw
rawreads.fasta
~~~
3. *FALCON*: all parameters were set to the defaults. Command line:

~~~
fc_run.py fc_run.cfg &>fc_run.out
~~~
4. *GPM*: manual edit with merging *de novo* assemblies from *MECAT2*, *Canu1.5*, and *FALCON*.

#### Polishing

1. *arrow*: all parameters were set to the defaults except `alignment length = 500 bp`. The *arrow* polish was carried out by the SMRT Link v6.0 webpage (https://www.pacb.com/support/software-downloads/).
2. *pilon1.18*: all parameters were set to the defaults and *pilon* polish was carried out as recommended at the SMRT Link v6.0 (https://www.pacb.com/support/software-downloads/).

#### BUSCO

The BUSCO3.0 version was employed in this study. Command line: `run_BUSCO.py-i genome.fasta -o genome -l embryophyta_odb9 -m genome -c 16`

#### RepeatMasker

The repeat sequences were employed with the library rice7.0.0_liban in-house. Command line: `RepeatMasker -pa 24 -x -no_is -nolow -cutoff 250 -lib rice7.0.0.liban.txt genome.fasta`

## Acknowledgements

This research was supported by the AXA Research Fund (International Rice Research Institute), the King Abdullah University of Science & Technology, and the Bud Antle Endowed Chair for Excellent in Agriculture (University of Arizona) to R.A.W., the Start-up Fund of Huazhong Agricultural University to J.Z., and funding from the Taiwan Council of Agriculture to IRRI. The BUSCO analysis data for maize was kindly provided by Dr. Wu and Dr. Li from the Institute of Plant Physiology and Ecology, and Dr. Wang from Shanghai Jiao Tong University. One of two TE libraries used for repeat analysis was provided by Dr. Eric Laserre (University of Perpignan, France)

## Author contributions

J.Z., K.M., D.C., M.L., N.A., N.R.S.H., H.L., R.M, and R.A.W. designed and conceived the research. D.C. and K.M. perform the population structure analysis. K.M., M.L., L.J.A., N.L. generated and provided SSD seed 12 *O. sativa* accessions. D.K., S.L., S.R., N.M prepared DNA and performed PacBio and Illumina sequencing. C.S.-S. managed all PacBio and Illumina sequence data processing and transfer. P.P. and V.L. generated all Bionano optical maps. J.Z. and Y.Z. performed sequence assembly. Y.Z. carried out genome size estimation, GPM editing, assembly polishing and data submission. V.L. and Y.Z. analyzed the Bionano optical maps and the validation of 12 PSRefSeqs. A.Z. and Y.Z. carried out TE prediction and structural analysis. Y.Z., N.A., A.Z., J.Z., D.C., M.L., K.M., N.M. and R.A.W. wrote and edited the paper. All authors read and approved the final manuscript.

## Supplementary Information

### Supplementary file1

**Supplementary Table 1**. Summary of Illumina genome survey sequences for 12 *Oryza sativa* genomes.

**Supplementary Table 2**. Genome features of *de novo* assemblies for 12 *Oryza sativa* accessions by Canu1.5, FALCON and MECAT2.

**Supplementary Table 3**. Genome features of 12 *Oryza sativa* accessions by GPM editing.

**Supplementary Table 4**. Chromosome length (Mb) of 12 *Oryza sativa* genomes.

**Supplementary Table 5**. Bionano optical map statistics of 12 *Oryza sativa* genomes.

**Supplementary Table 6**. Summary of large structural variation (>50 bp) by comparison of each of 16 genomes to every other genome (including 12 genomes from this study and 4 previously reported: MH63, ZS97, N 22 and the IRGSP RefSeq).

### Supplementary file2

**Supplementary Figure 1**. Admixture results for K=5 to 15. The samples are grouped according to the new classification. At K=9,12,13, the Q matrices converged to two different modes, differing according to whether ind1A is split, or tropical japonica.

**Supplementary Figure 2**. Length distribution of PacBio long reads used for 12 *Oryza sativa* genome assemblies.

**Supplementary Figure 3**. K-mer analysis of Illumina short sequences that were used for genome size estimation with the GCE program.

**Supplementary Figure 4**. Bionano Access visualization view for 12 *de novo* assemblies with Bionano optical maps and their underlying alignments.

**Supplementary Figure 5**. Summary of missing genes in the BUSCO gene space evaluation of 12 *de novo Oryza sativa* assemblies, 4 public *Oryza sativa* PSRefSeqs and 3 high-quality *Zea mays* genomes.

