## Supplementary_file2 for "Twelve Platinum-Standard Reference Genomes Sequences (PSRefSeq) that complete the full range of genetic diversity of Asian rice"

**Supplementary files**

**Supplementary file 1**

Supplementary Table 1. Summary of Illumina genome survey sequences for 12 *Oryza sativa* genomes.
Supplementary Table 2. Genome features of *de novo* assemblies for 12 *Oryza sativa* accessions by Canu1.5, FALCON and MECAT2.
Supplementary Table 3. Genome features of 12 *Oryza sativa* accessions by GPM editing.
Supplementary Table 4. Chromosome length (Mb) of 12 *Oryza sativa* genomes.
Supplementary Table 5. Bionano optical map statistics of 12 *Oryza sativa* genomes.
Supplementary Table 6. Summary of large structural variation (>50 bp) by comparison of each of 16 genomes to every other genome (including 12 genomes in this study and 4 previously reported: MH63, ZS97, N 22 and the IRGSP RefSeq).

**Supplementary file2**

**Supplementary Figure 1**. Admixture results for K=5 to 15. The samples are grouped according to the new classification. At K=9,12,13, the Q matrices converged to two different modes, differing according to whether ind1A is split, or tropical japonica.
**Supplementary Figure 2**. Length distribution of PacBio long reads used for 12 *Oryza sativa* genome assemblies.
**Supplementary Figure 3**. K-mer analysis of Illumina short sequences that were used for genome size estimation with the GCE program.
**Supplementary Figure 4**. Bionano Access visualization view for 12 *de novo* assemblies with Bionano optical maps and their underlying alignments.
**Supplementary Figure 5**. Summary of missing genes in the BUSCO gene space evaluation of 12 *de novo Oryza sativa* assemblies, 4 public *Oryza sativa* PSRefSeqs and 3 high-quality *Zea mays* genomes.

Supplementary Figure 1

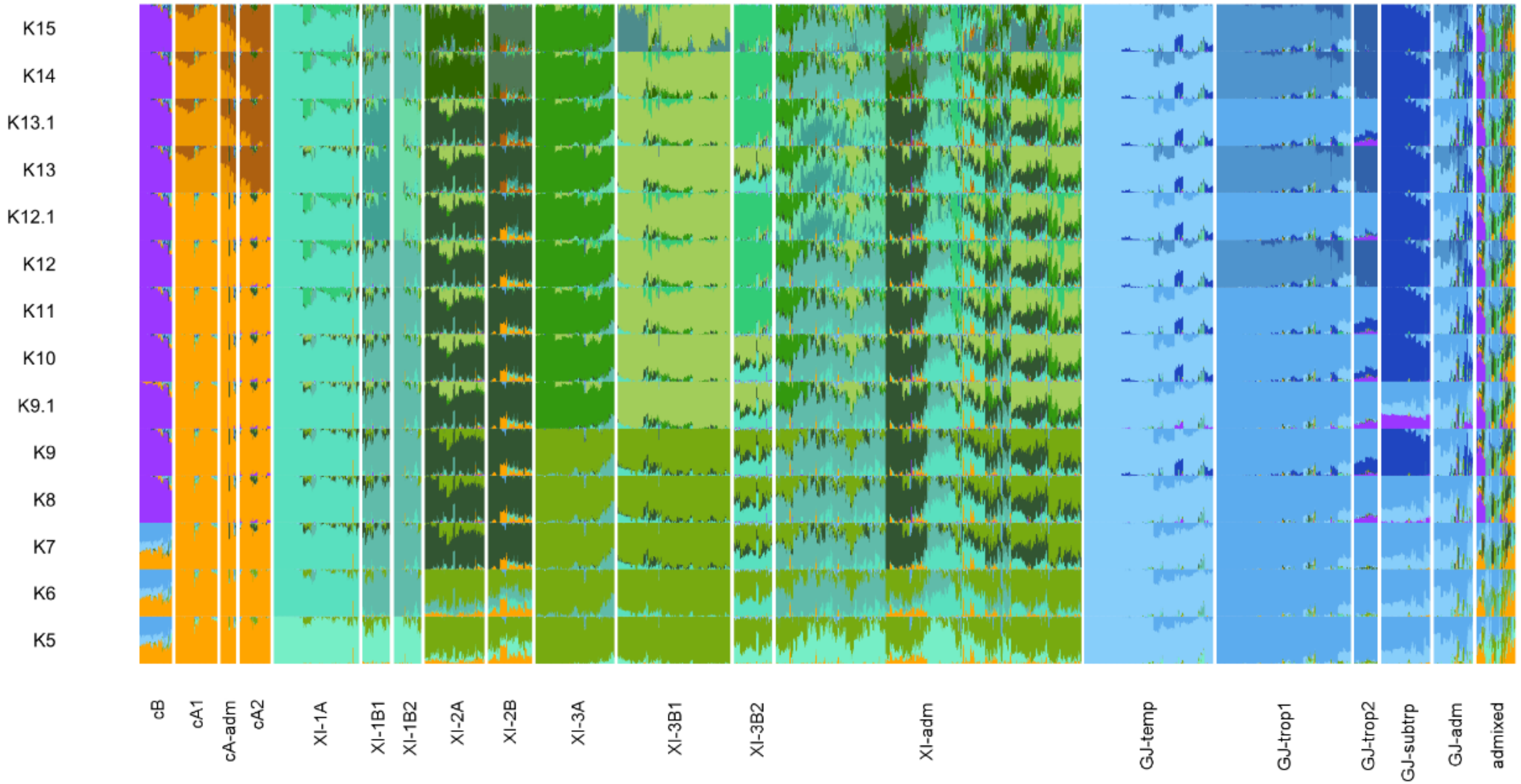

Supplementary Figure 2

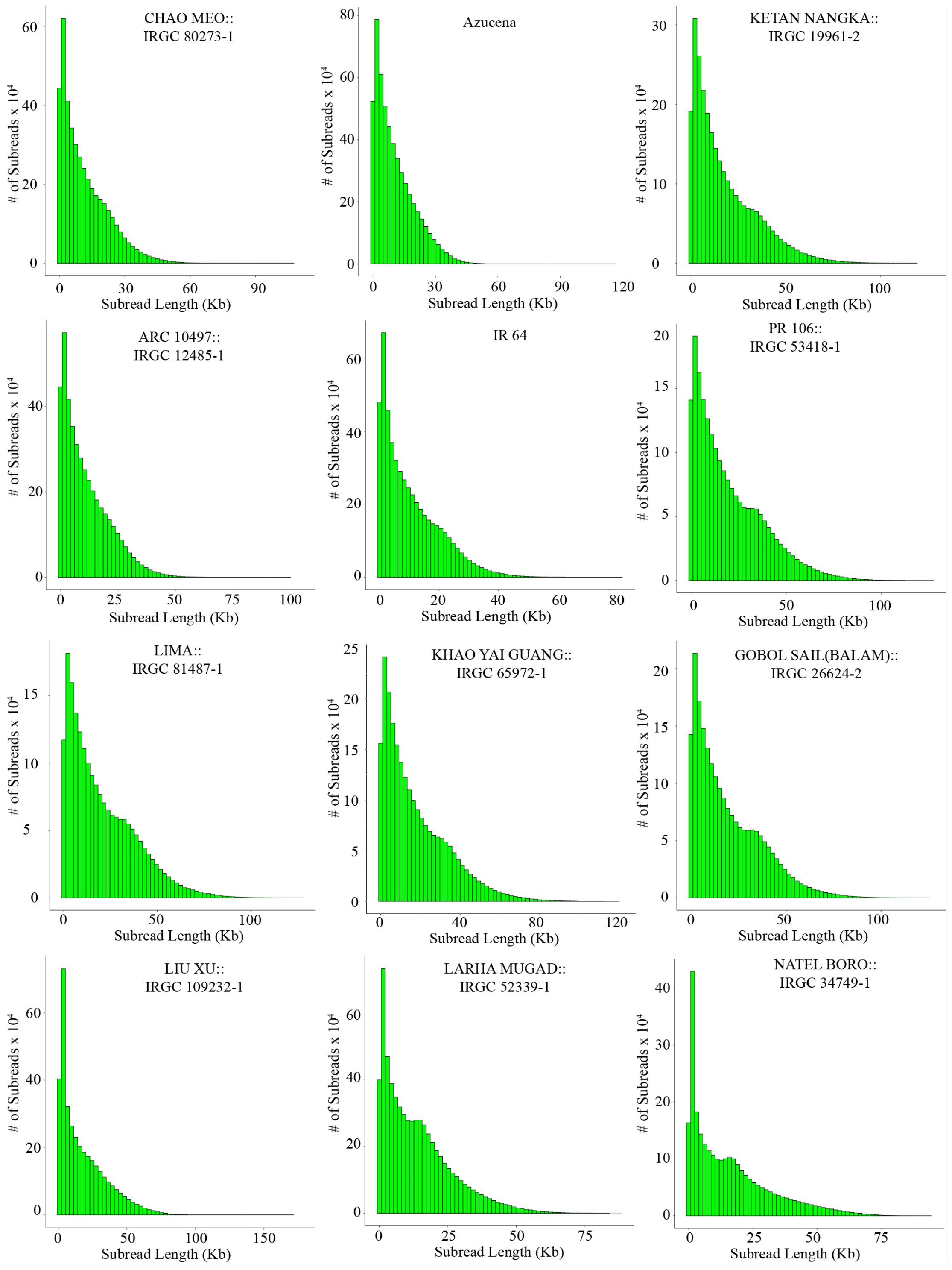

### Supplementary Figure S3

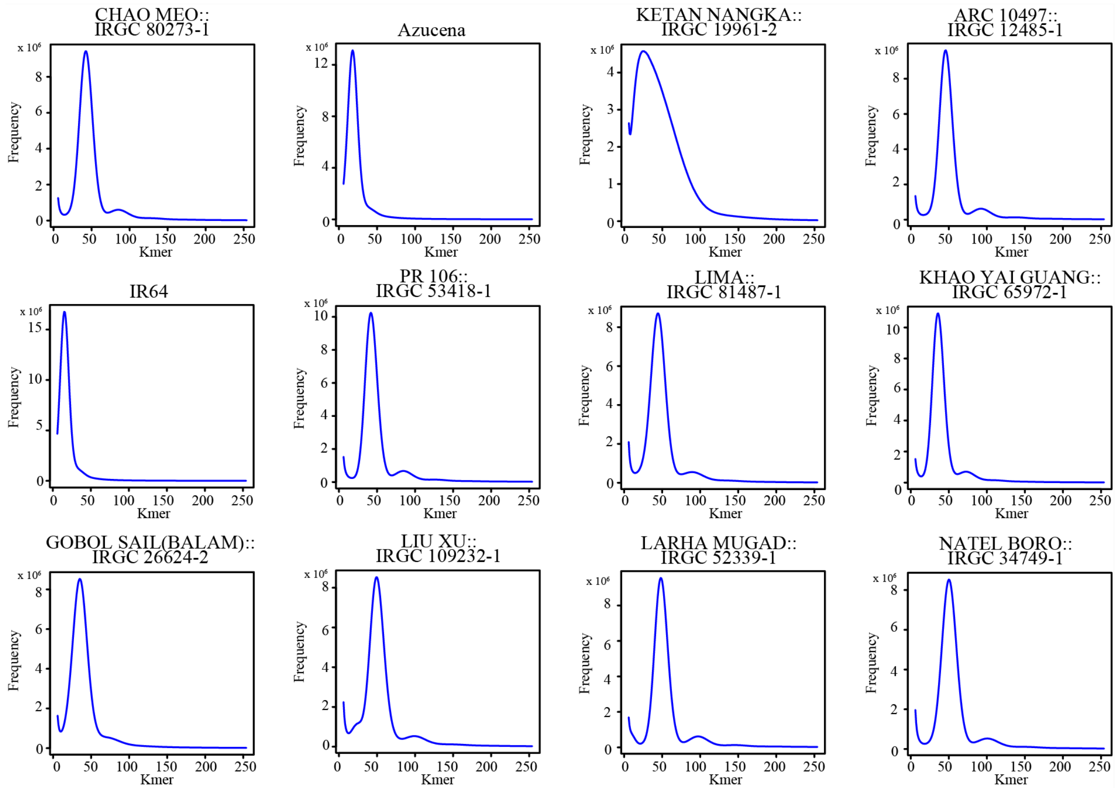

Supplementary Figure 4

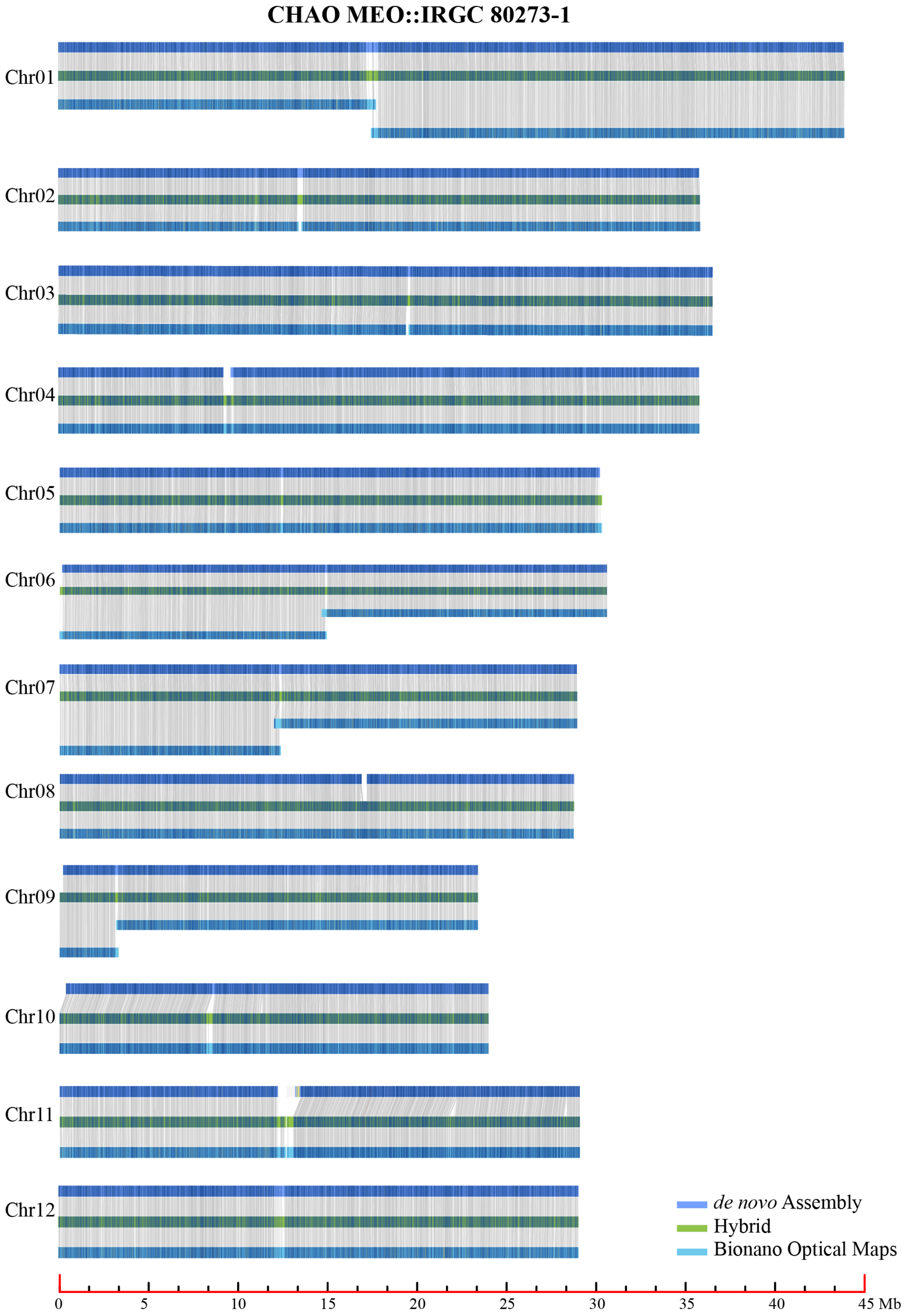

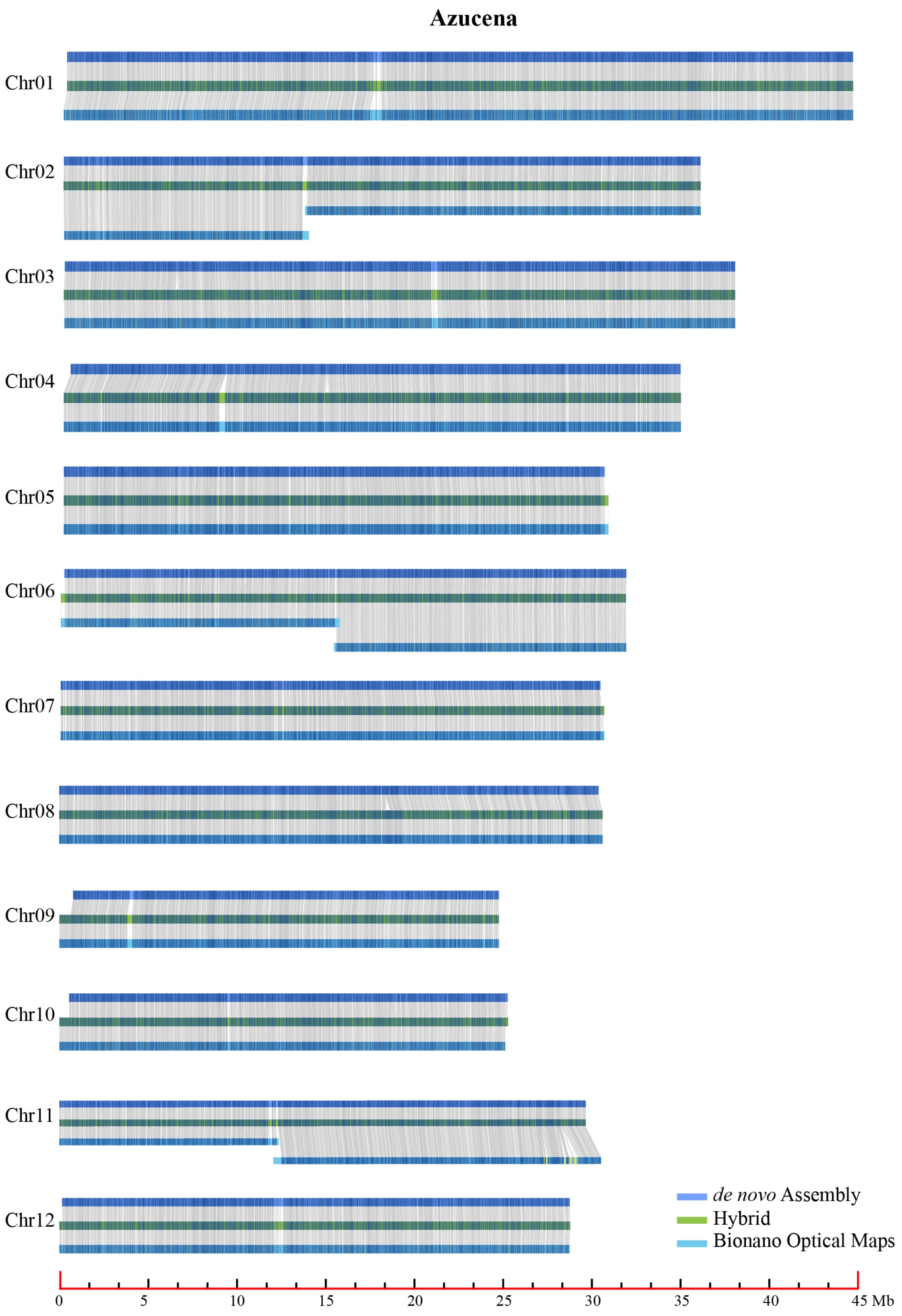
Azucena
Chr01
Chr02
Chr03
Chr04
Chr05
Chr06
Chr07
Chr08
Chr09
Chr10
Chr11
Chr12
de novo Assembly
Hybrid
Bionano Optical Maps
0
5
10
15
20
25
30
35
40
45 Mb

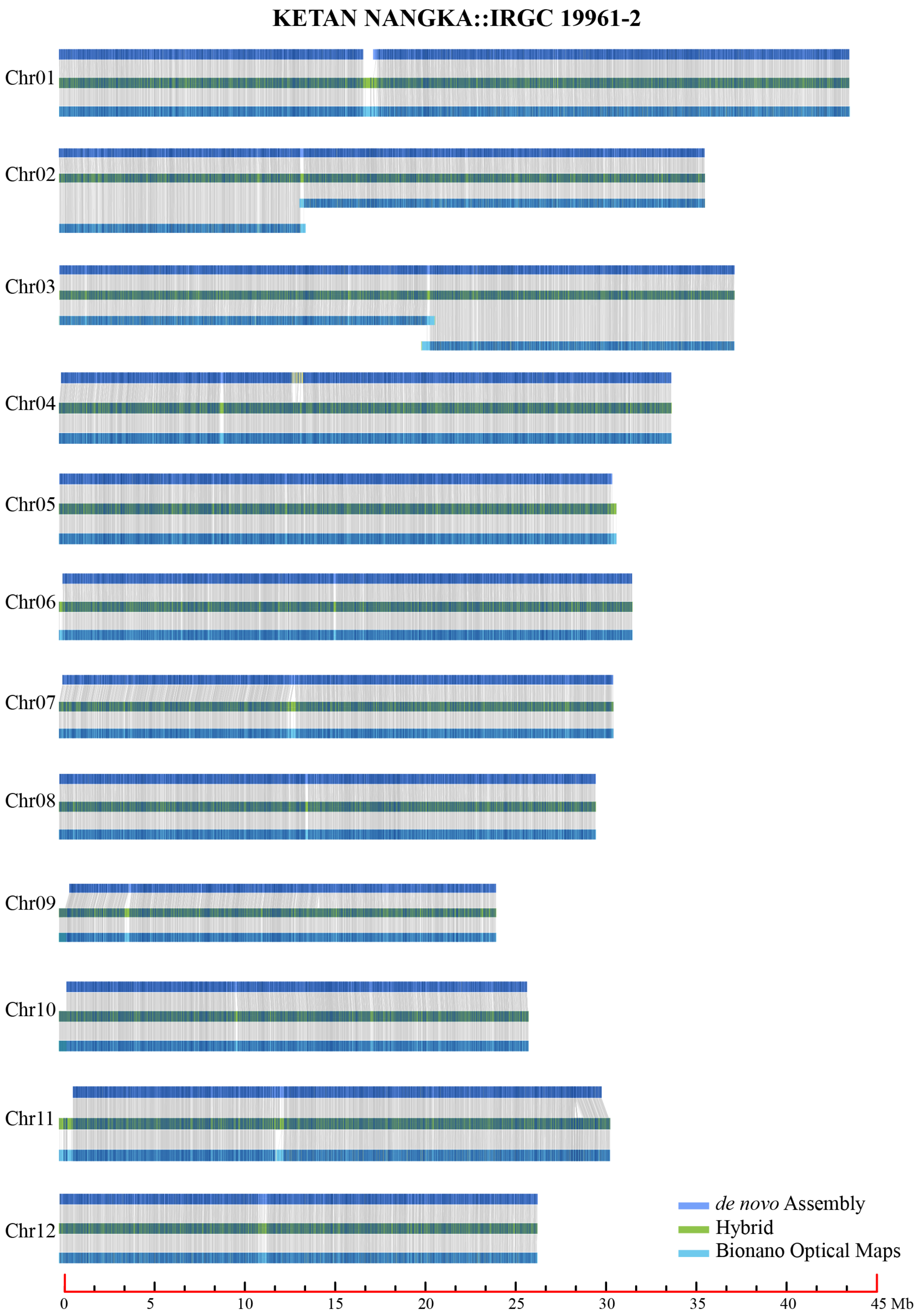
KETAN NANGKA::IRGC 19961-2
Chr01
Chr02
Chr03
Chr04
Chr05
Chr06
Chr07
Chr08
Chr09
Chr10
Chr11
Chr12
de novo Assembly
Hybrid
Bionano Optical Maps
0
5
10
15
20
25
30
35
40
45 Mb

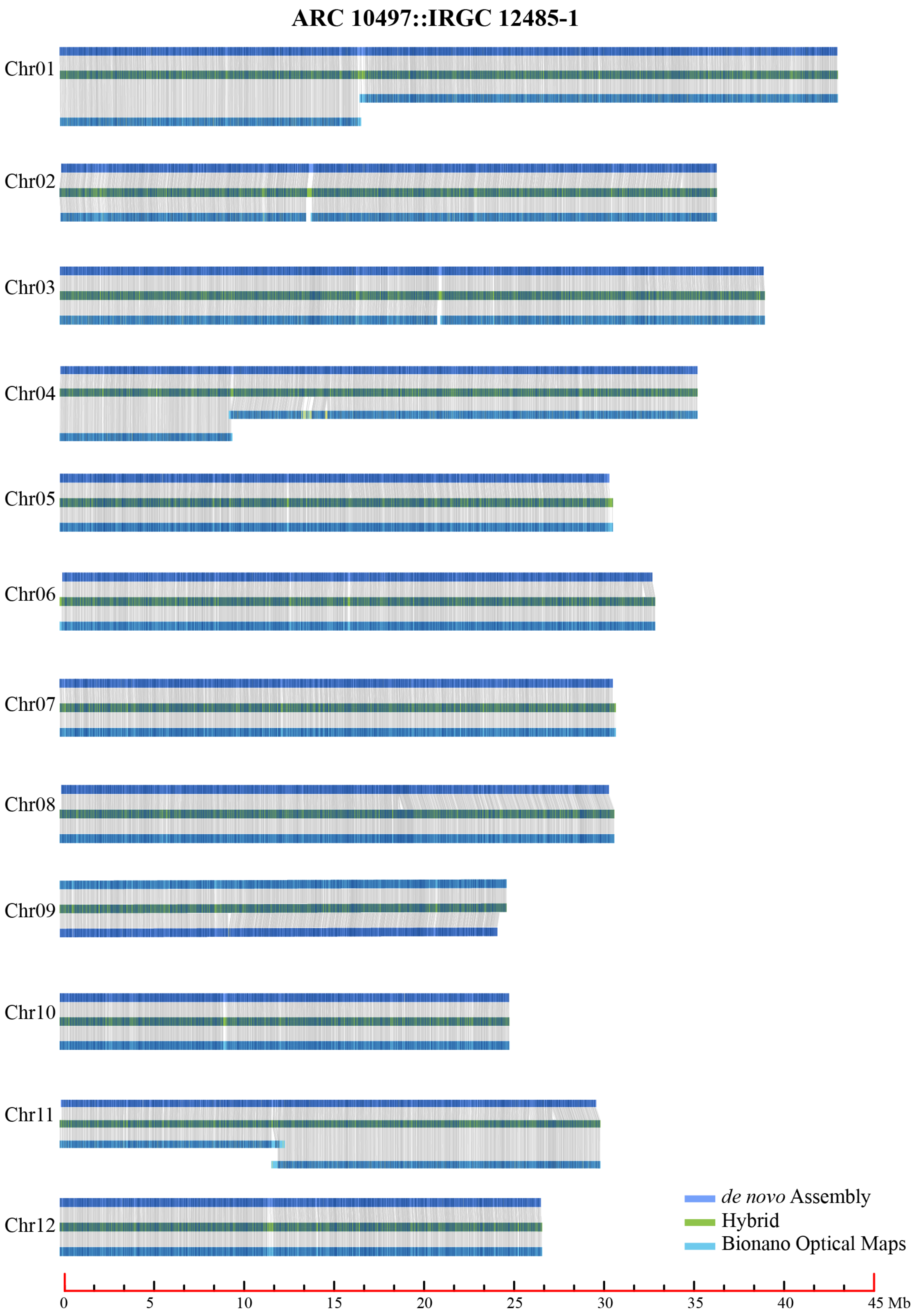
ARC 10497::IRGC 12485-1
Chr01
Chr02
Chr03
Chr04
Chr05
Chr06
Chr07
Chr08
Chr09
Chr10
Chr11
Chr12
de novo Assembly
Hybrid
Bionano Optical Maps
0
5
10
15
20
25
30
35
40
45 Mb

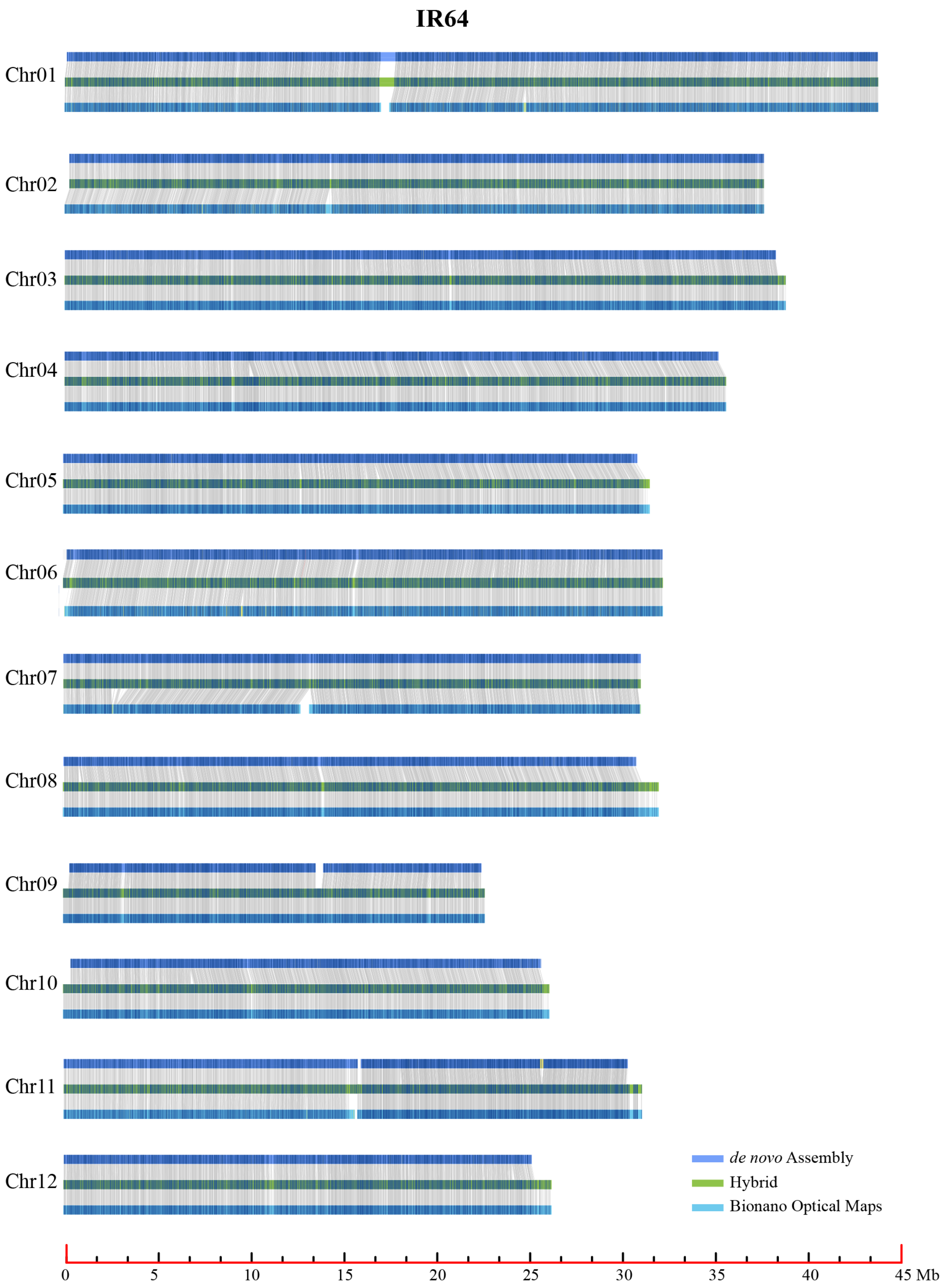

IR64
Chr01
Chr02
Chr03
Chr04
Chr05
Chr06
Chr07
Chr08
Chr09
Chr10
Chr11
Chr12
de novo Assembly
Hybrid
Bionano Optical Maps
0
5
10
15
20
25
30
35
40
45 Mb

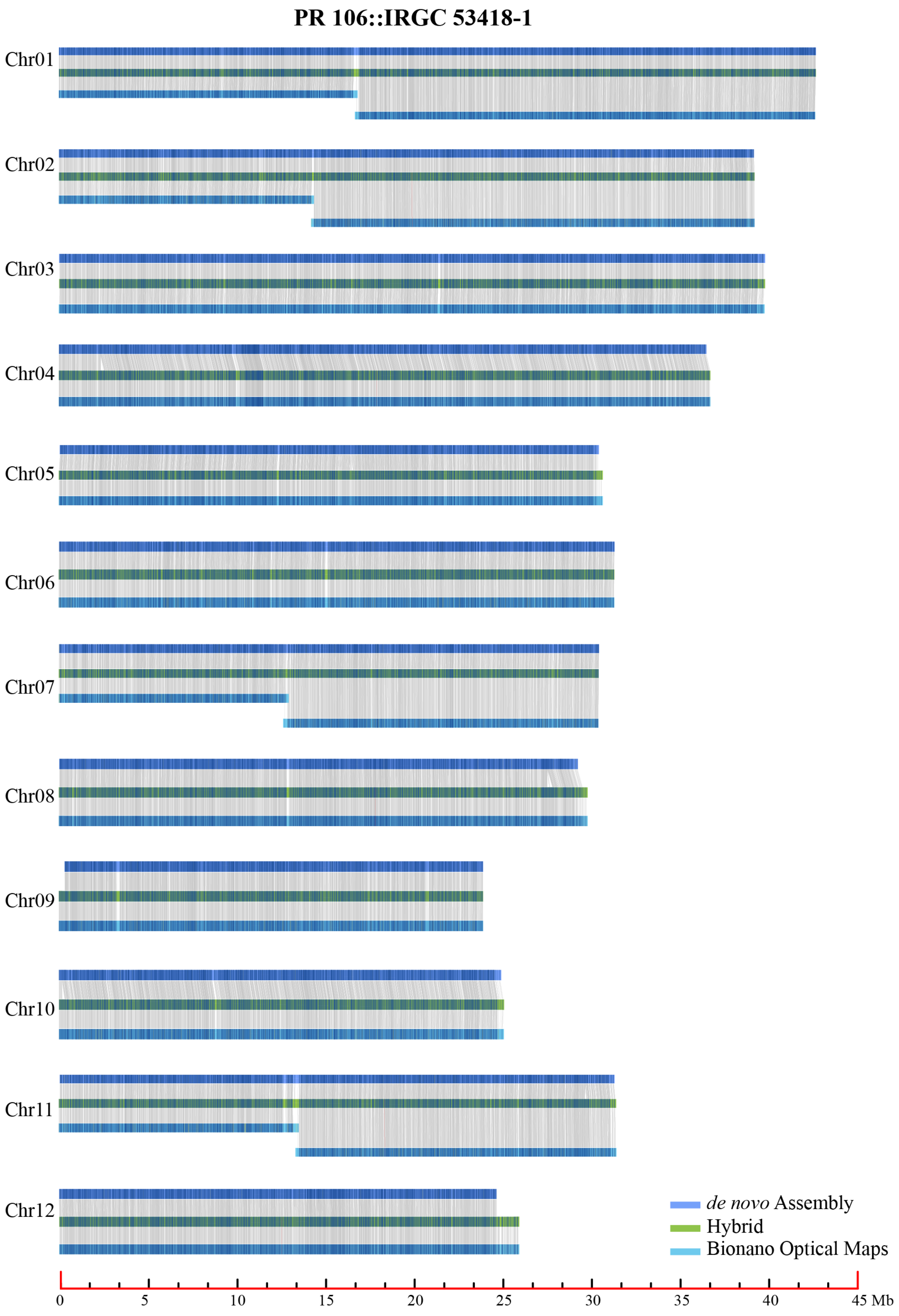
PR 106::IRGC 53418-1
Chr01
Chr02
Chr03
Chr04
Chr05
Chr06
Chr07
Chr08
Chr09
Chr10
Chr11
Chr12
de novo Assembly
Hybrid
Bionano Optical Maps
0
5
10
15
20
25
30
35
40
45 Mb

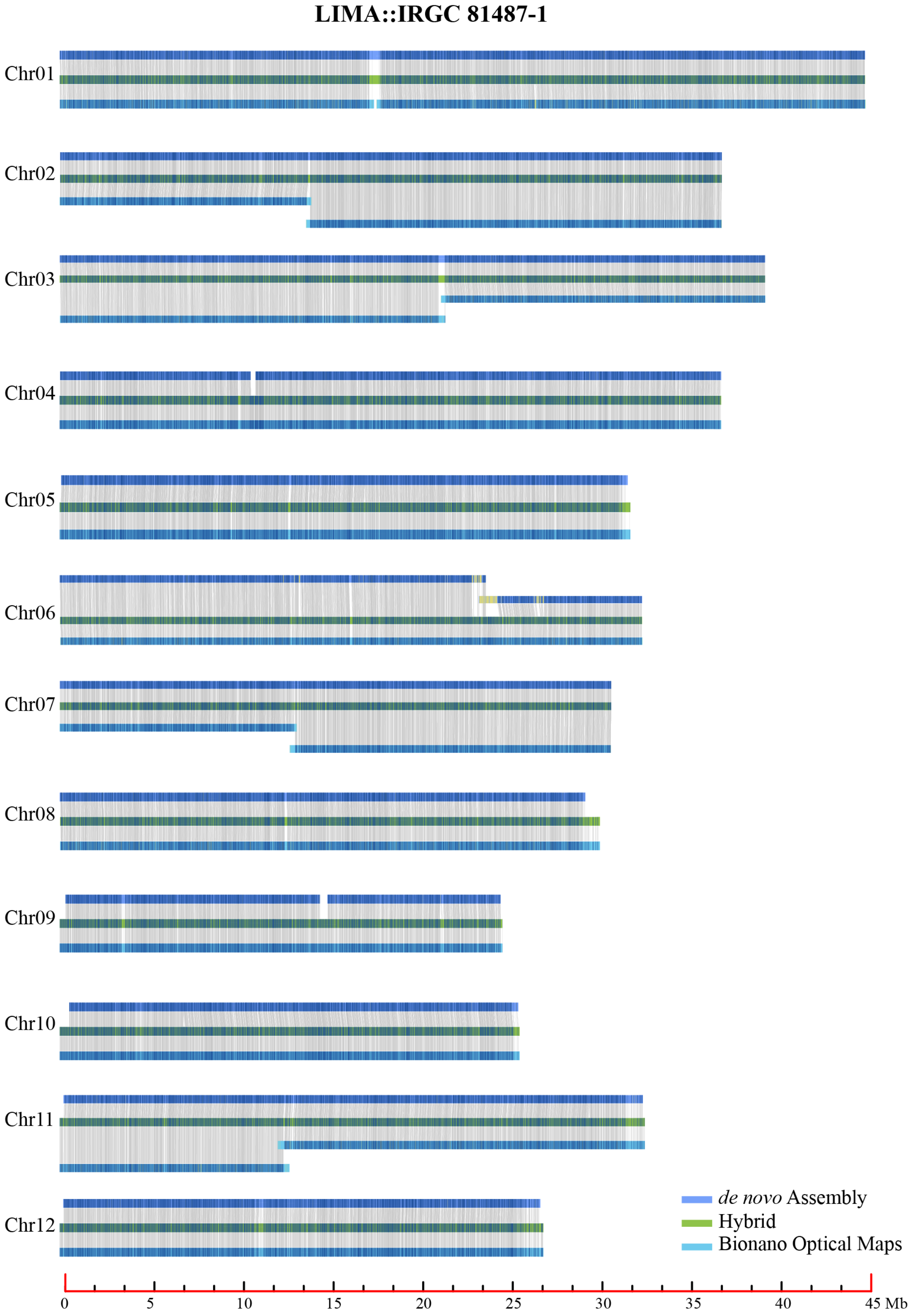

LIMA::IRGC 81487-1
Chr01
Chr02
Chr03
Chr04
Chr05
Chr06
Chr07
Chr08
Chr09
Chr10
Chr11
Chr12
de novo Assembly
Hybrid
Bionano Optical Maps
0
5
10
15
20
25
30
35
40
45 Mb

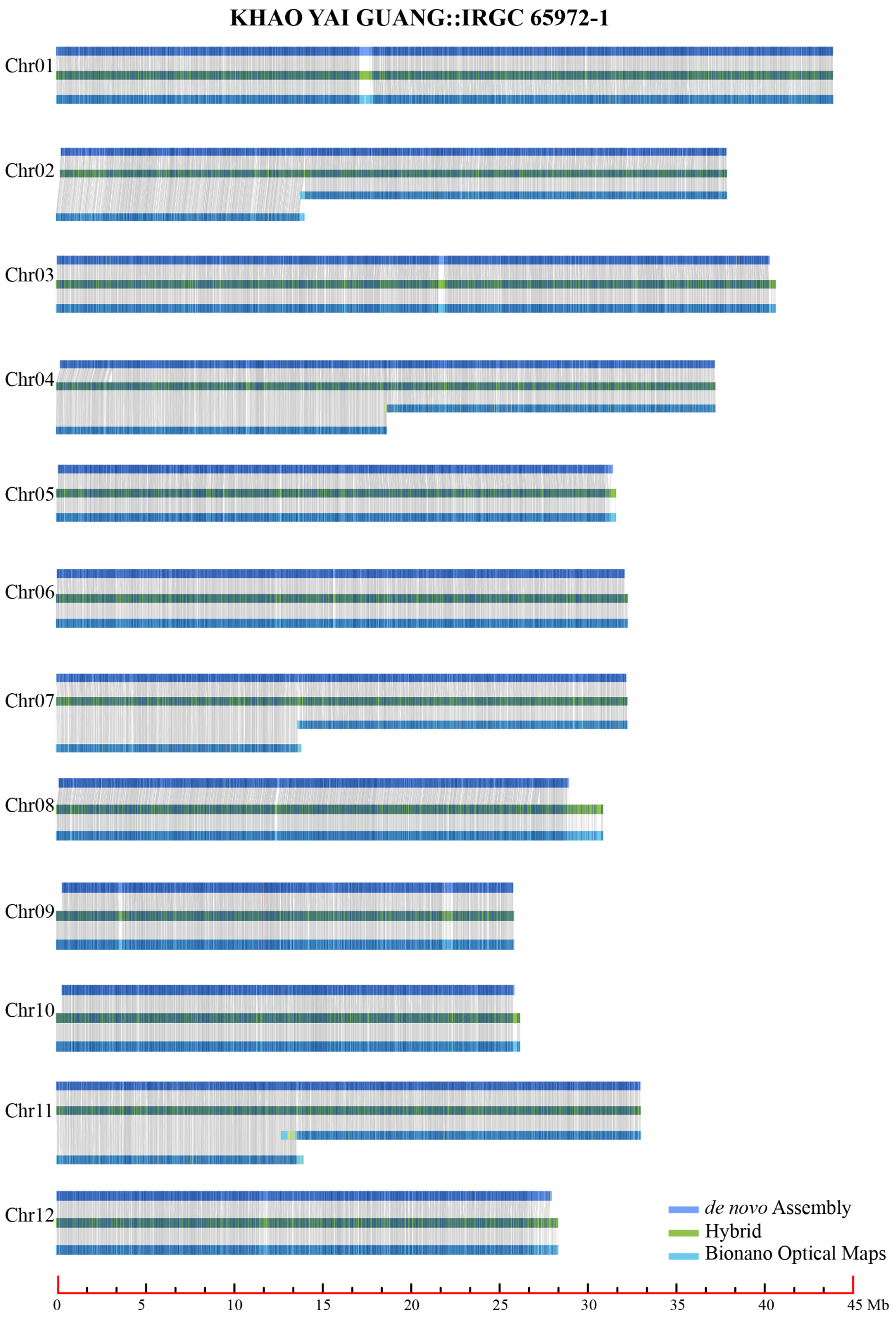
KHAO YAI GUANG::IRGC 65972-1
Chr01
Chr02
Chr03
Chr04
Chr05
Chr06
Chr07
Chr08
Chr09
Chr10
Chr11
Chr12
de novo Assembly
Hybrid
Bionano Optical Maps
0
5
10
15
20
25
30
35
40
45 Mb

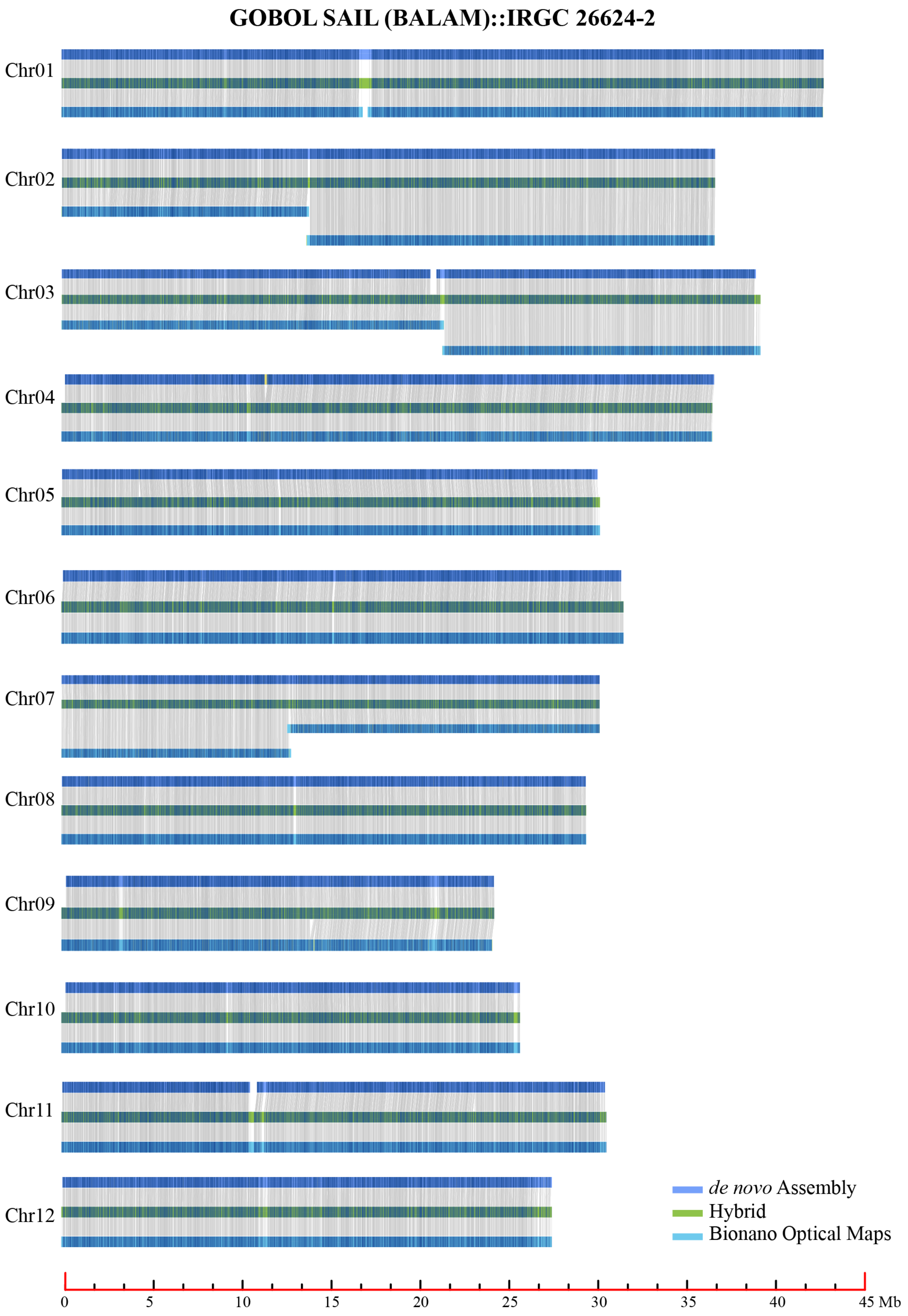
GOBOL SAIL (BALAM)::IRGC 26624-2
Chr01
Chr02
Chr03
Chr04
Chr05
Chr06
Chr07
Chr08
Chr09
Chr10
Chr11
Chr12
de novo Assembly
Hybrid
Bionano Optical Maps
0
5
10
15
20
25
30
35
40
45 Mb

#### LIU XU::IRGC 109232-1

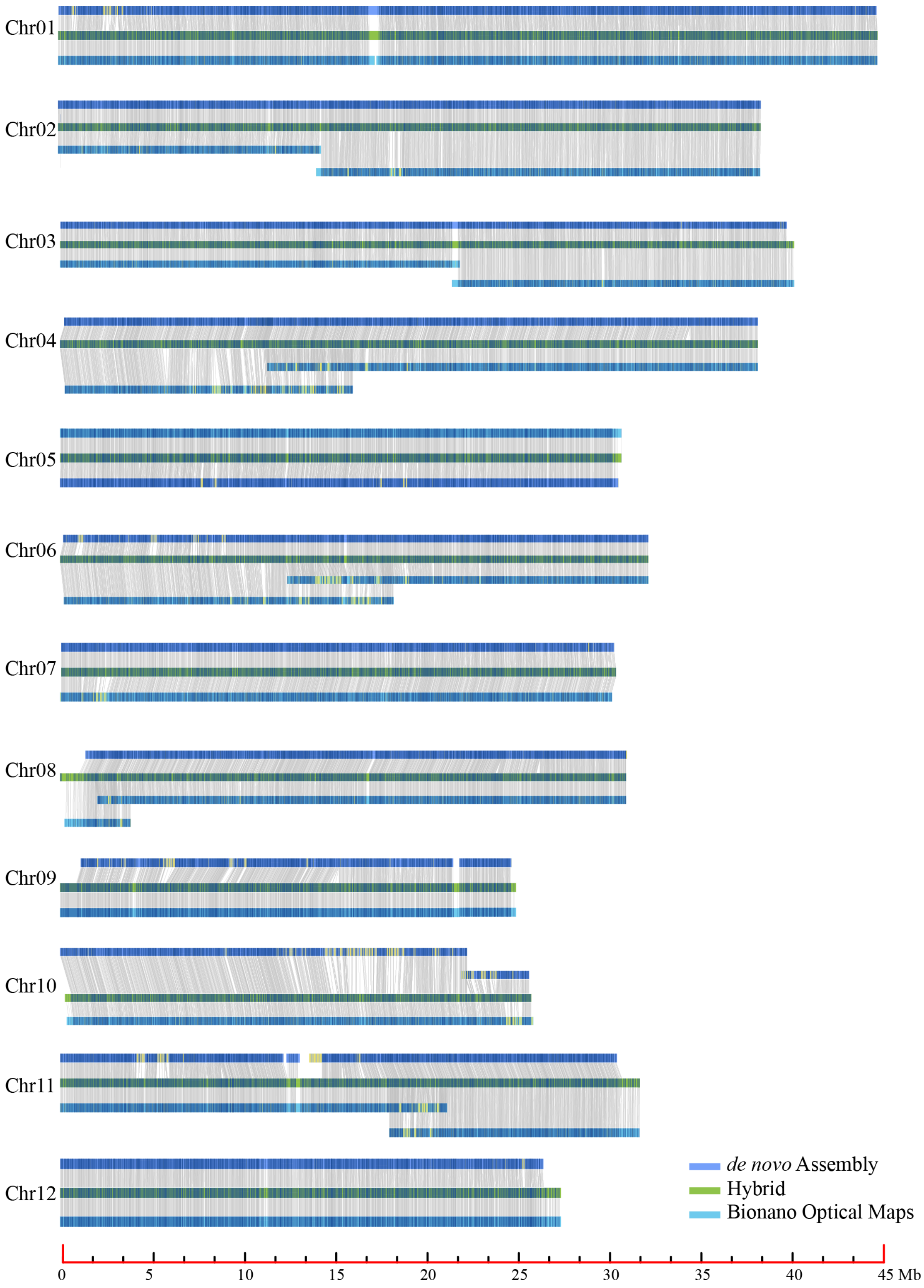

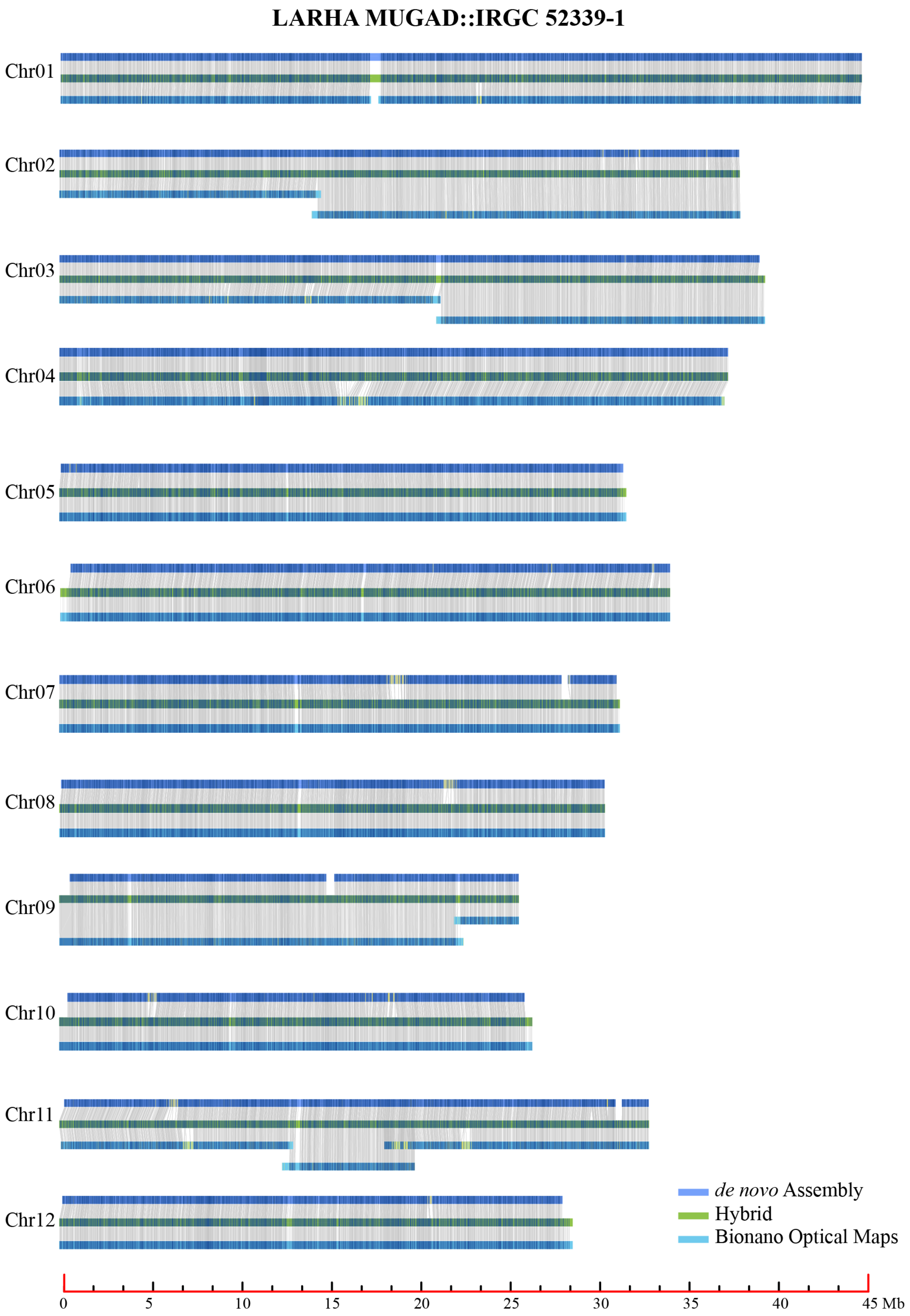
LARHA MUGAD::IRGC 52339-1
Chr01
Chr02
Chr03
Chr04
Chr05
Chr06
Chr07
Chr08
Chr09
Chr10
Chr11
Chr12
de novo Assembly
Hybrid
Bionano Optical Maps
0
5
10
15
20
25
30
35
40
45 Mb

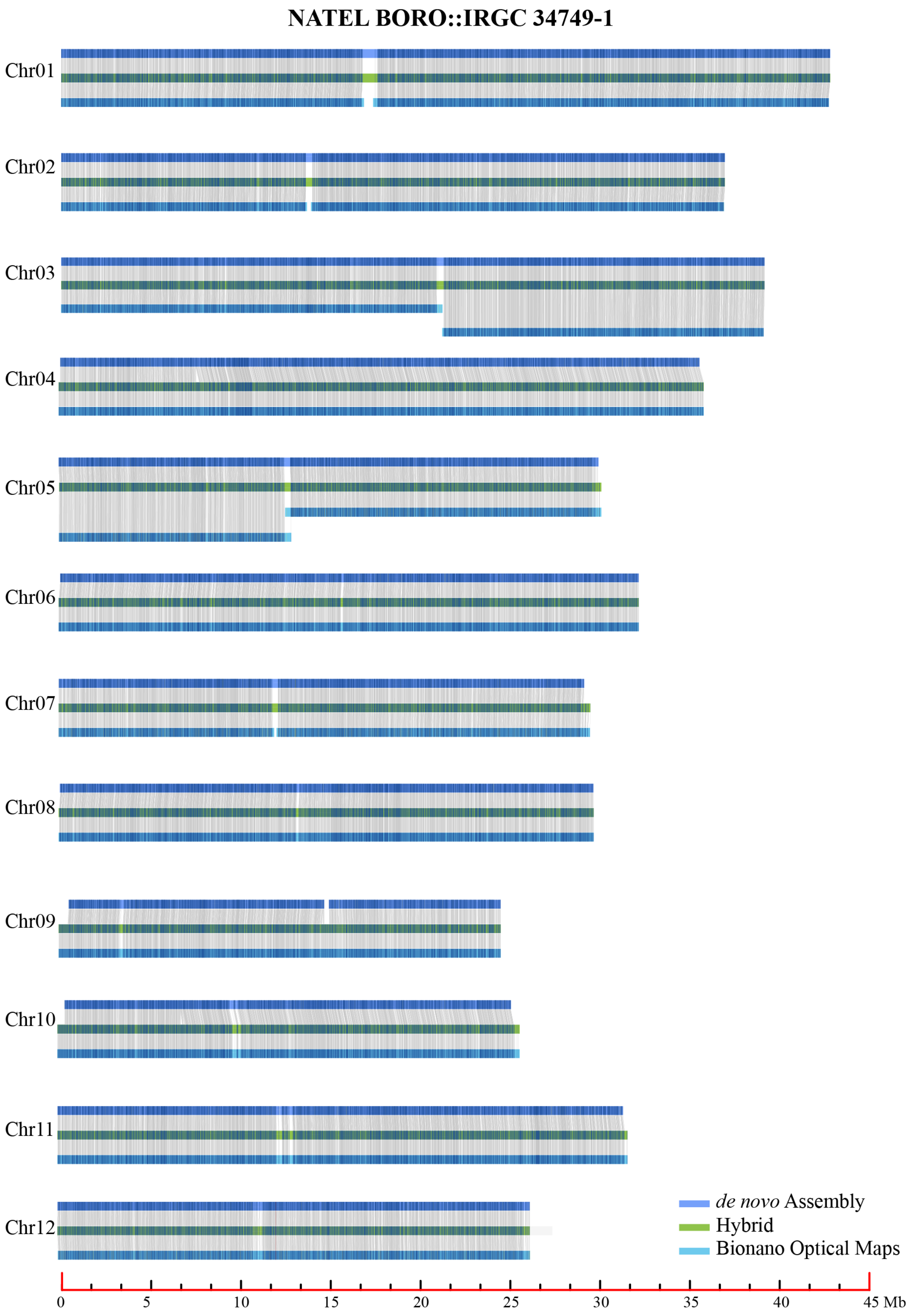

NATEL BORO::IRGC 34749-1
Chr01
Chr02
Chr03
Chr04
Chr05
Chr06
Chr07
Chr08
Chr09
Chr10
Chr11
Chr12
de novo Assembly
Hybrid
Bionano Optical Maps
0
5
10
15
20
25
30
35
40
45 Mb

Supplementary Figure 5

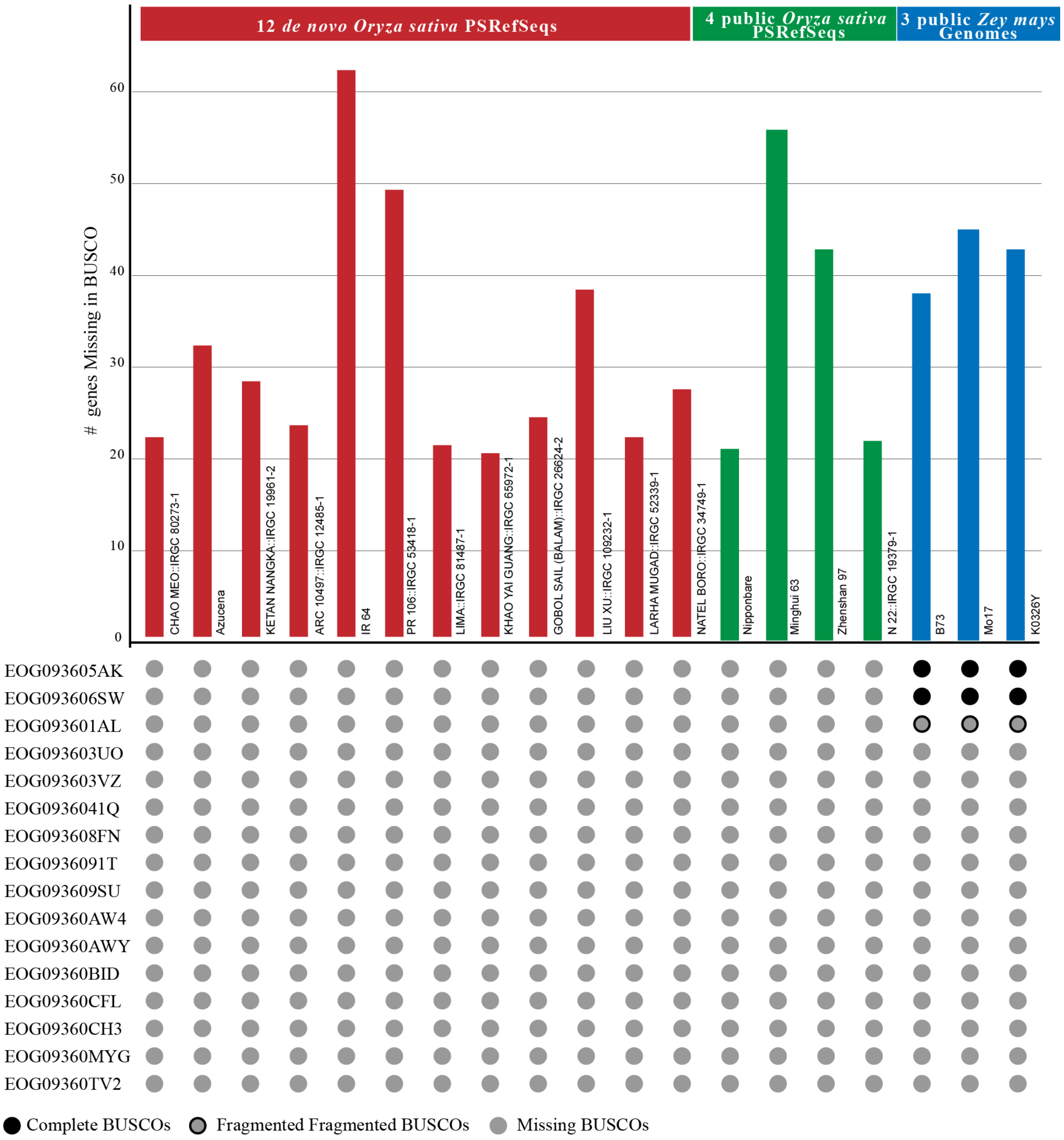
